# Taxonomic variability over functional stability in the microbiome of Cystic Fibrosis patients chronically infected by *Pseudomonas aeruginosa*

**DOI:** 10.1101/609057

**Authors:** Giovanni Bacci, Giovanni Taccetti, Daniela Dolce, Federica Armanini, Nicola Segata, Francesca Di Cesare, Vincenzina Lucidi, Ersilia Fiscarelli, Patrizia Morelli, Rosaria Casciaro, Anna Negroni, Alessio Mengoni, Annamaria Bevivino

**Affiliations:** Department of Biology, University of Florence, Sesto Fiorentino, 50019, Florence, Italy; Cystic Fibrosis Center, Anna Meyer Children’s University Hospital, Department of Pediatrics Medicine, Florence, 50139, Italy; Centre for Integrative Biology, University of Trento, Trento, 38122, Italy; Children’s Hospital and Research Institute Bambino Gesù, Rome, 00165, Italy; Cystic Fibrosis Center, IRCCS G. Gaslini Institute, Department of Pediatrics, Genoa, 16147, Italy; Department for Sustainability, Italian National Agency for New Technologies, Energy and Sustainable Economic Development, ENEA, 00123 Rome, Italy

**Keywords:** cystic fibrosis, lung microbiome, longitudinal studies, metagenome composition, antibiotic resistance genes

## Abstract

Although the cystic fibrosis (CF) lung microbiome has been characterized in several studies, little is still known about the functions harboured by those bacteria, and how they change with disease status and antibiotic treatment. The aim of this study was to investigate the taxonomic and functional temporal dynamics of airways microbiome in a cohort of CF patients. Multiple sputum samples were collected over 15 months from 22 patients with chronic *P. aeruginosa* infection, for a total of 79 samples. DNA extracted from samples was subjected to shotgun metagenomic sequencing allowing either strain-level taxonomic profiling and assessment of the functional metagenomic repertoire. High inter-patient taxonomic heterogeneity was found with short-term compositional changes during exacerbations and following antibiotic treatment. Each patient exhibited distinct sputum microbial communities at the taxonomic level, and strain-specific colonization of traditional CF pathogens, including *P. aeruginosa*, and emerging pathogens. Sputum microbiome was found to be extraordinarily resilient following antibiotic treatment, with rapid recovery of taxa and metagenome-associated gene functions. In particular, a large core set of genes, including antibiotic resistance genes, were shared across patients despite observed differences in clinical status or antibiotic treatment, and constantly detected in the lung microbiome of all subjects independently from known antibiotic exposure, suggesting an overall microbiome-associated functions stability despite taxonomic fluctuations of the communities.

**IMPORTANCE:** While the dynamics of CF sputum microbial composition were highly patient-specific, the overall sputum metagenome composition was stable, showing a high resilience along time and antibiotic exposure. The high degree of redundancy in the CF lung microbiome could testifies ecological aspects connected to the disease that were never considered so far, as the large core-set of genes shared between patients despite observed differences in clinical status or antibiotic treatment. Investigations on the actual functionality (e.g. by metatranscriptomics) of the identified core-set of genes could provide clues on genetic function of the microbiome to be targeted in future therapeutic treatments.

## Introduction

Bacterial lung infections reduce life expectancy in most individuals with cystic fibrosis (CF) (1). Sputum bacterial loads remain equally high both during periods of clinical stability and during pulmonary exacerbations (2), the latter of which contribute to the irreversible decline of lung function. Though much is known about microbes that cause respiratory infections in CF (3), how microbes contribute to exacerbations is still poorly understood. In the past years, studies employing DNA-based analyses of the airway microbiota of CF patients have reported somewhat discordant results. Indeed, while some studies showed a largely stable airway microbiota during clinical change and antibiotic treatment (4), other did not, suggesting changes the involvement of some microbial taxa in exacerbation (5, 6). Most of these used 16S rRNA gene sequencing, yielding the identities and relative abundances of the taxa present (i.e., the microbiota), but without providing any strain-level or functional (meaning based on functional genes) information (i.e., the metagenome) (7). These latter characteristics are particularly relevant for studying host-microbiome interactions. Indeed, defining the dynamics of individual microbial strains provides important information regarding how specific sub-lineages of pathogens persist and relate to clinical change. On the other hand, studying the microbial genetic repertoire, e.g. antibiotic resistance and virulence-related genes, with respect to clinical status or treatment can identify mechanisms of microbial persistence and pathogenesis (8–10). Until now, few longitudinal studies, with a limited number of patients, focusing only on CF airway microbiota have been performed (11–13). Moreover, studies on the complete CF microbiome (microbiota and metagenome) are few and on a limited number of patients (13–16) or focused on specific metabolic functions (17). In this work, we studied the temporal dynamics of CF sputum microbiomes, focusing on patients with moderate-severe lung disease, chronically infected by *Pseudomonas aeruginosa*. In a previous work (18), chronic infection with *P. aeruginosa* has been found to be associated with dysbiosis in the lungs of patients with CF. The authors suggested that the dominance of one species remodels the lung microbiota and may promote severity of CF lung disease. A more detailed taxonomic and functional analysis could help elucidating the mechanisms leading to chronic infection with *P. aeruginosa* and the microbial factors that contribute to the global changes of their lung microbiome. In the present study, a shotgun metagenomic approach was used (19) to detect the entire sputum microbial genomic repertoire down to the strain level (20, 21). A cohort of 22 patients with moderate-severe lung disease, grouped according to homozygosity versus heterozygosity for ΔF508 (also known as F508del) in the CFTR gene and chronically infected with *P. aeruginosa*, was selected and followed over 15 months during which 8 patients underwent exacerbation events. We aimed to determine the composition of sputum microbiomes for these patients when longitudinally sampled during periods of stability and exacerbation, defining the relationship between clinical status, sputum microbial metabolic gene repertoire, and the antibiotic-resistance (AR) gene composition of sputum bacterial community, providing a previously unknown, high-resolution view of CF sputum microbiome dynamics.

## Results

### Patients and sampling

Twenty-two patients with CF were enrolled (15 females and seven males) who had moderate-severe lung disease (30 < %FEV_1_ < 70) and were chronically infected by *P. aeruginosa*, according to the Leeds criteria (22). During the study period, patients were treated with maintenance antibiotics (aerosol) and only a subset (n = 8) received clinical additional antibiotics (oral or/and intravenous) for a pulmonary exacerbation (CFPE) (Table 1 and supplementary materials Table S1). Among the 22 subjects, 8 were diagnosed with exacerbations during the study period. In total, 79 sputum samples were collected and analyzed using shotgun metagenomic sequencing.

**TABLE 1.** Characteristics of patients enrolled in the study

| ID | Hospital | Genotype | Gender | FEV<sub>1</sub><br>status | Age | n | EX | %FEV<sub>1</sub> |
| --- | --- | --- | --- | --- | --- | --- | --- | --- |
| B01 | OPBG | ΔF508/2183AA->G | M | S | 27 | 5 | yes | 37.0 ± 1.70 |
| B02 | OPBG | ΔF508/N1303K | F | SD | 26 | 3 | no | 54.7 ± 3.48 |
| B03 | OPBG | ΔF508/4016insT | F | S | 30 | 4 | no | 55.0 ± 1.08 |
| B06 | OPBG | ΔF508/ΔF508 | F | SD | 21 | 4 | no | 60.2 ± 3.42 |
| G10 | Gaslini | ΔF508/ΔF508 | M | S | 51 | 4 | no | 54.0 ± 3.08 |
| G24 | Gaslini | ΔF508/ΔF508 | F | S | 49 | 3 | yes | NA ± NA |
| G28 | Gaslini | ΔF508/ΔF508 | F | NA | 38 | 2 | no | 42.5 ± 1.50 |
| G30 | Gaslini | ΔF508/ΔF508 | F | S | 50 | 1 | no | 54 |
| G31 | Gaslini | G1244E/G42X | F | SD | 53 | 2 | no | 41.5 ± 1.50 |
| G34 | Gaslini | ΔF508/ΔF508 | F | S | 39 | 1 | no | 47 |
| M05 | Meyer | ΔF508/ΔF508 | M | SD | 32 | 4 | no | 34.8 ± 0.85 |
| M19 | Meyer | ΔF508/ΔF508 | M | S | 24 | 4 | no | 44.0 ± 2.04 |
| M21 | Meyer | ΔF508/N1303K | M | SD | 27 | 4 | yes | 51.5 ± 4.35 |
| M22 | Meyer | ΔF508/2789+5G->A | F | S | 29 | 5 | yes | 50.4 ± 1.03 |
| M23 | Meyer | ΔF508/G542X | F | S | 30 | 4 | yes | 37.0 ± 1.47 |
| M24 | Meyer | ΔF508/ΔF508 | M | S | 32 | 4 | no | 35.2 ± 0.85 |
| M25 | Meyer | ΔF508/296+1G->T | F | SD | 41 | 4 | no | 42.5 ± 2.02 |
| M26 | Meyer | ΔF508/3849+10 | F | SD | 49 | 5 | yes | 39.6 ± 1.94 |
| M28 | Meyer | ΔF508/N1303K | M | S | 23 | 4 | no | 39.0 ± 1.08 |
| M29 | Meyer | ΔF508/G542X | F | S | 12 | 4 | no | 43.5 ± 3.75 |
| M31 | Meyer | ΔF508/ΔF508 | F | SD | 11 | 3 | yes | 32.7 ± 4.41 |
| M33 | Meyer | ΔF508/G85E | F | SD | 13 | 5 | yes | 35.4 ± 5.78 |
| **Total:**<br>**22** | Gaslini:6<br>Meyer:12<br>OPBG:4 | Heterozygote :47<br>Homozygote :29<br>Other:2 | F:15<br>M:7 | S:12<br>SD:9 | 32.1 ± 2.73 | 79 | no:14<br>yes:8 | 43.5 ± 1.09 |

ID, study id; Hospital, hospital in which patient has been enrolled [OPBG=Children's Hospital and Research Institute Bambino Gesù (Rome, Italy); Gaslini=G. Gaslini Institute (University of Genoa, Genoa, Italy); Meyer=Cystic Fibrosis Center, Anna Meyer Children's University Hospital (Florence, Italy)]; Genotype, CFTR genotype; Gender, gender; Age, enrollment's age; n, number of samples collected; EX, yes if an exacerbation event has occurred during the study (no otherwise) (2); FEV<sub>1</sub>, mean value of forced expiratory volume in 1 second plus/minus the standard error on the mean; heterozygote and homozygote refers to ΔF508 genotype; %FEV<sub>1</sub> status: S = with a rate decline lower than 1.5%, SD = with a rate decline higher than 5% (44).

### Airway microbiomes are taxonomically distinct and show patient-specific strain colonization

The overall taxonomic representation of the microbiomes from the 79 samples is reported in Fig. 1a and 1b, whereas a summary of obtained reads per sample was reported in supplementary materials Table S2. Firmicutes, Proteobacteria, Bacteroidetes, and Actinobacteria were the most represented phyla. A high relative abundance of the “classical” CF bacterial signatures (taxa), such as *Staphylococcus aureus* and *Pseudomonas aeruginosa*, and non-traditional CF taxa, such as *Rothia mucilaginosa*, and *Prevotella melaninogenica* (all present in the top-10 species within each phylum, Fig. 1b), was found. These species, indeed, represent the 49% of all detected taxa as reported in supplementary materials Table S3.

**FIGURE 1.**
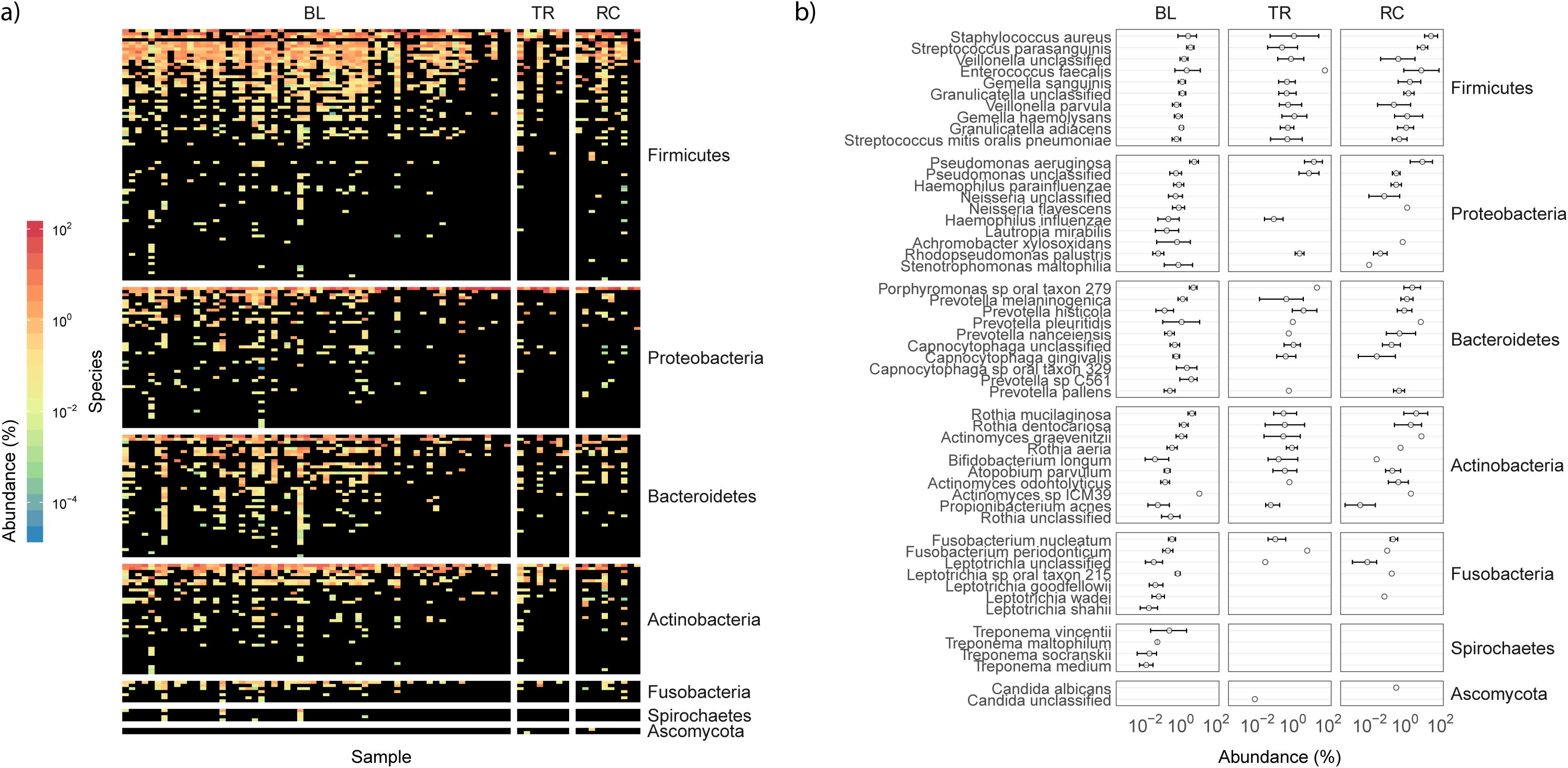
Taxonomic distribution in patients enrolled in the study. a) The taxonomic distribution of all species detected using MetaPhlAn2 was reported in each row of the matrix whereas columns represent samples collected during the study. Colors from dark blue to red were used to report “copies per million” (CPM) values as obtained from HUMAnN2 with black reporting a CPM value of zero. The plot was divided according to patient status: BL, baseline; TR, treatment; RC, recovery. Species were ordered according to their mean abundance and grouped according to their Phylum. b) The mean abundance value of the top-ten species (if available) detected within each Phylum was reported together with the standard error. The relative abundance of taxa is reported (Abundance %).

Although principal coordinates analysis showed that subjects did not cluster based on treatment events and/or genotype (Fig. 2a), the PERMANOVA analysis (Table 2) reported a significant effect of both factors. The R^2^ values, namely the proportion of variance explained by the factor considered, were very low (Table 2, R^2^ = 0.03 for both factors, p-values < 0.05) probably due to intra-patient heterogeneity. CFTR genotype did not influence the effect of antibiotic treatment (viz. exacerbation) on sputum microbiota, nor vice versa (p-value > 0.05, treatment-genotype interaction effect). Subject effect was predominant with an R^2^ value of 0.52. A high fraction (more than 50%) of the total variance can be thus explained by inter-subject variation. Neither FEV_1_ nor time showed any significant relationship with taxonomy or functional profile (Table 2).

**FIGURE 2.**
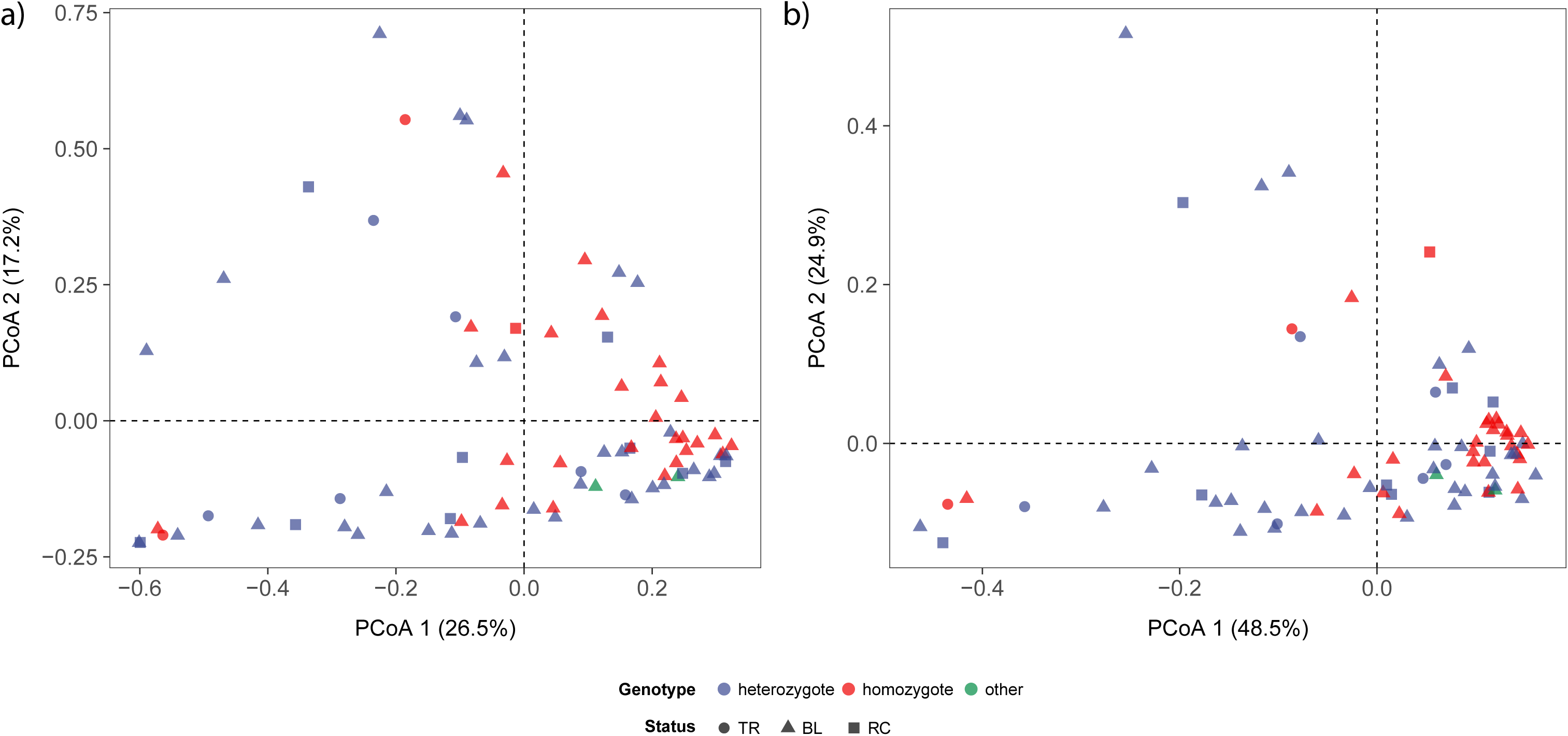
Ordination analyses based on a) taxonomic assignments and b) pathway distribution detected with MetaPhlAn2 and HUMAnN2, respectively. Ordination analyses were conducted using the Bray-Curtis dissimilarity index and ordered following the principle coordinate decomposition method (PCoA). The percentage of variance explained by each coordinate was reported between round brackets. Homozygote and heterozygote refer to ΔF508 mutation of CFTR gene. BL, baseline; TR, treatment; RC, recovery.

**TABLE 2.**
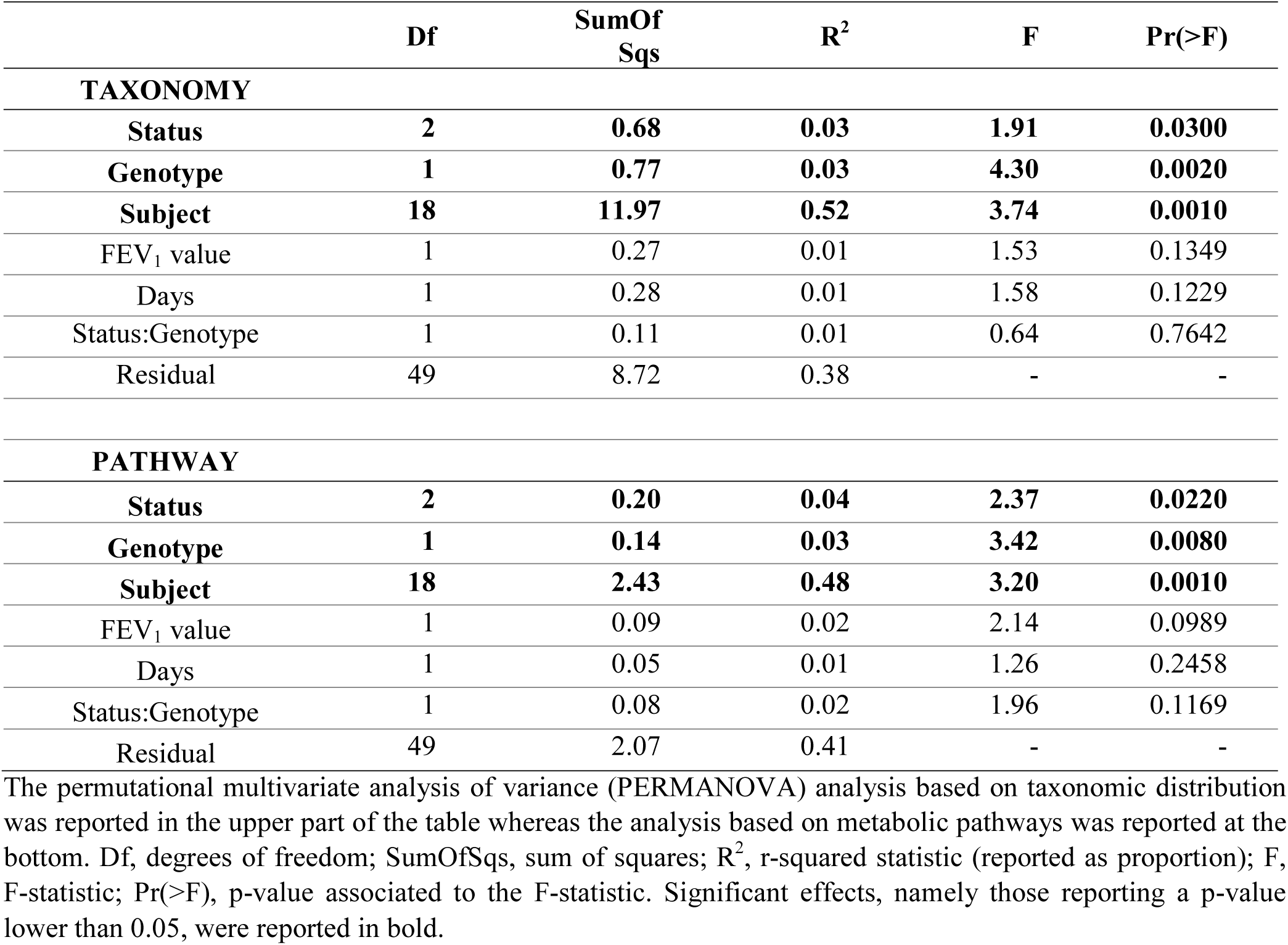
Permutational multivariate analysis of variance on both taxonomic distribution and metabolic pathways

We then performed a strain-level analysis of the sputum microbiomes. This analysis demonstrated, in samples from the same patient but at different time points, that bacterial lineages were in general, closely related and tightly clustered together, confirming a patient-specific bacterial colonization (Fig. 3 and supplementary materials Fig. S1).

**FIGURE 3.**
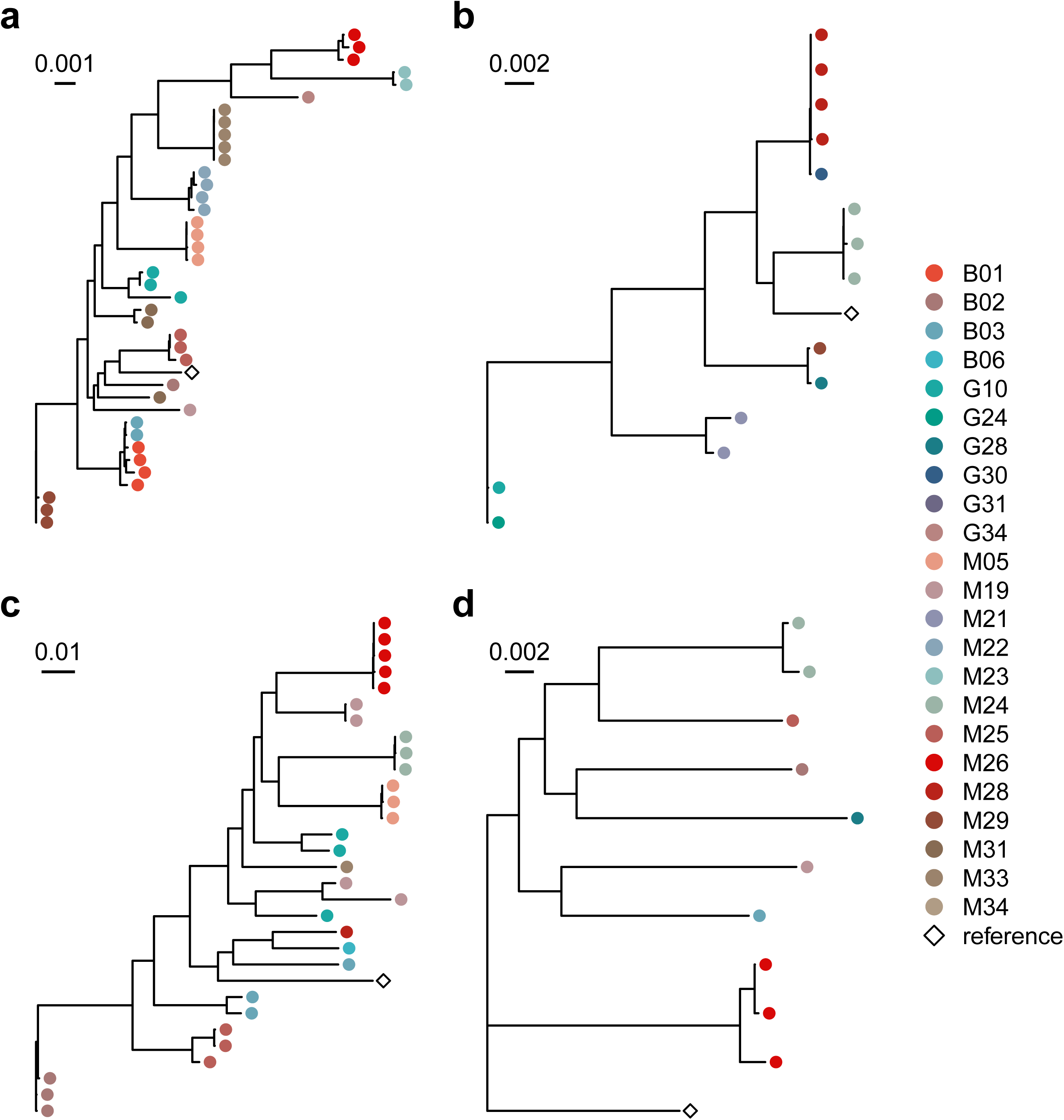
Strain-level phylogenetic trees of the main CF pathogens detected in the study. Phylogenetic trees obtained through StrainPhlAn pipeline were reported for the main pathogenic signatures of CF disease: a) *Pseudomonas aeruginosa*; b) *Staphylococcus aureus*; c) *Rothia mucilaginosa*; d) *Prevotella melaninogenica*. Points at the end of each clade are colored according to patients so that two points with the same color, in the same tree, represent the same species in two different time points, for the same patient.

Alpha diversity analyses were consistent with the results above. Bacterial diversity measures (Shannon and inverse Simpson indices) varied according to clinical status, genotype, and subject (Fig. 4a, supplementary materials Fig. S2 and Table S4). Samples collected during clinical treatments exhibited lower microbial diversity than samples collected at either baseline or recovery visits, highlighting the role of clinical treatments in perturbing CF lung communities (supplementary materials Table S5).

**FIGURE 4.**
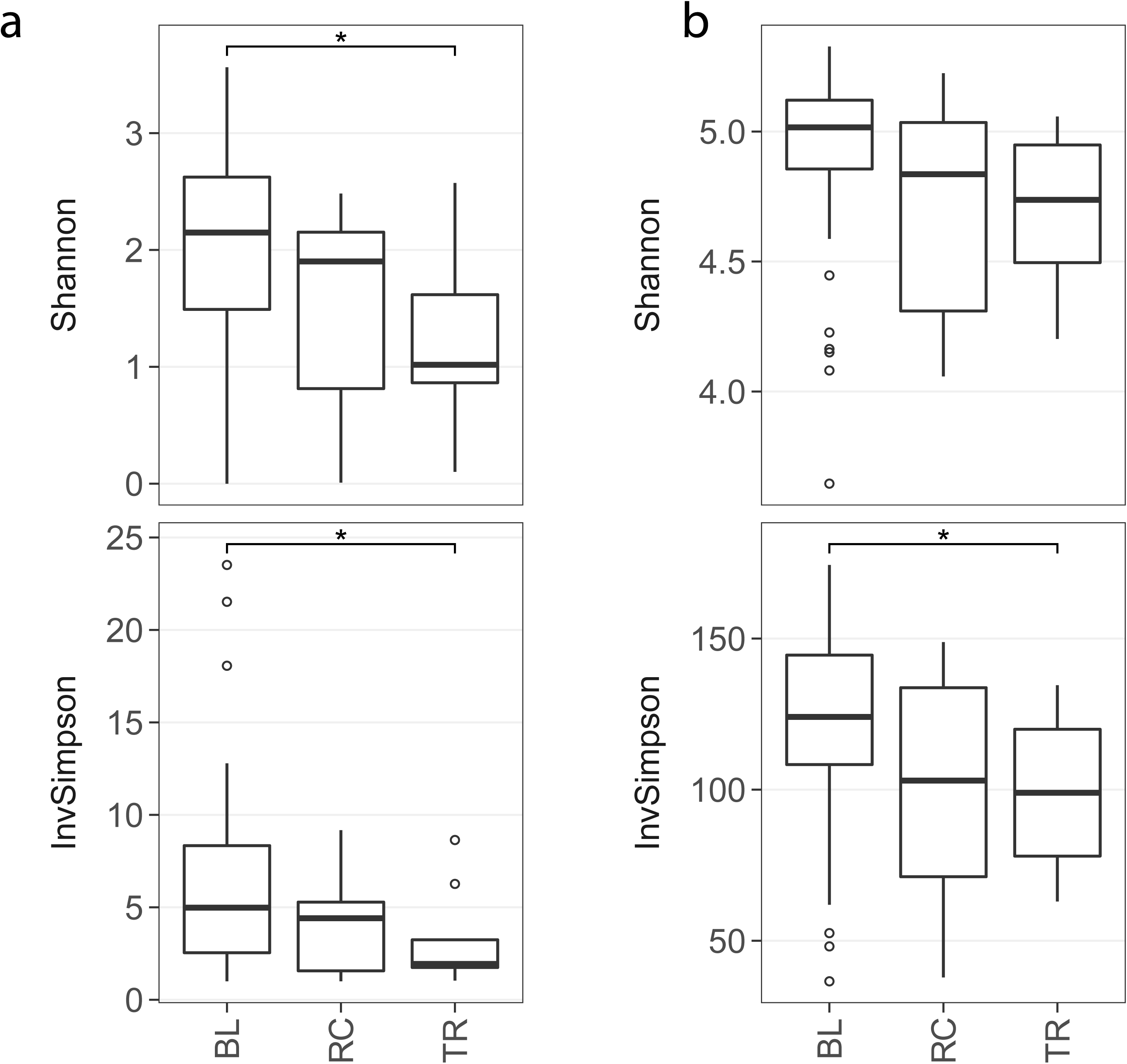
Differences across exacerbation events. The effect of an exacerbation event on alpha diversity was inspected using both the Shannon index and the inverse Simpson index. Diversity indexes were computed for both a) taxonomic signature and b) metabolic pathways. BL, baseline; TR, treatment; RC, recovery. Each box shows the “interquartile range” (IQR) that is the differences between the third and the first quartile of data (the 75^th^ and the 25^th^ percentile). Horizontal bars are medians whereas whiskers represent the minimum and maximum values defined as Q1 – (1.5 x IQR) and Q3 + (1.5 x IQR), respectively. Observations that fell outside minimum and maximum values were defined as outliers and reported using white points.

### Airway microbiomes are functionally consistent and show subject-specific distribution patterns

The results of functional metagenomic analyses were consistent with the taxonomic findings described above. Exacerbation events and patient genotype significantly impacted pathway distribution (Table 2, R^2^ values of 0.04 and 0.03 respectively, p-values < 0.05), though with less an effect than that of subject (R^2^ = 0.48). The sample distribution according to the ordination analysis (PCoA) was very heterogeneous with no sharp differences according to genotypes or exacerbation events. Alpha diversity dropped significantly in samples collected during exacerbation events, but the drop was significant only considering the inverse Simpson index (p-value = 0.036, Fig. 4b and supplementary materials Table S4). Overall, the pathway distribution was more consistent with respect to the taxonomic one, with biosynthetic pathways being the most represented functional category (Fig. 5, supplementary materials Fig. S2 and Table S5). Pathways were mainly detected in members of the phyla Firmicutes and Proteobacteria, followed by Bacteroidetes and Actinobacteria. Even if these results confirmed the results from the analysis of the taxonomic distribution, metabolic pathways showed a more consistent distribution across samples. Indeed, the beta-diversity analysis on both taxonomic and functional distribution showed a lower similarity based on taxonomy in respect with pathways (Supplementary materials Table S6, Fig. 6a and 6b). These results were additionally confirmed by the differential abundance analysis. For contrasts made within each genotype, only 40 pathways reported significant differences across exacerbation statuses (p-values < 0.05 and |log(fold-change)| > 5) all in the homozygote group (Supplementary materials Fig. S3 and Table S7), whereas, considering all samples together, no pathway was found to be more abundant in one condition in respect with another (data not shown). These results confirmed the extraordinary resilience of the CF microbiome evident at the taxonomic level is also exhibited from a functional perspective, indicating that neither clinical change nor antibiotic treatments are accompanied by changes in sputum microbial functions; for example, antibiotics do not appear to select for or against specific functions. To test this specifically, we focused on known antibiotic resistance genes.

**FIGURE 5.**
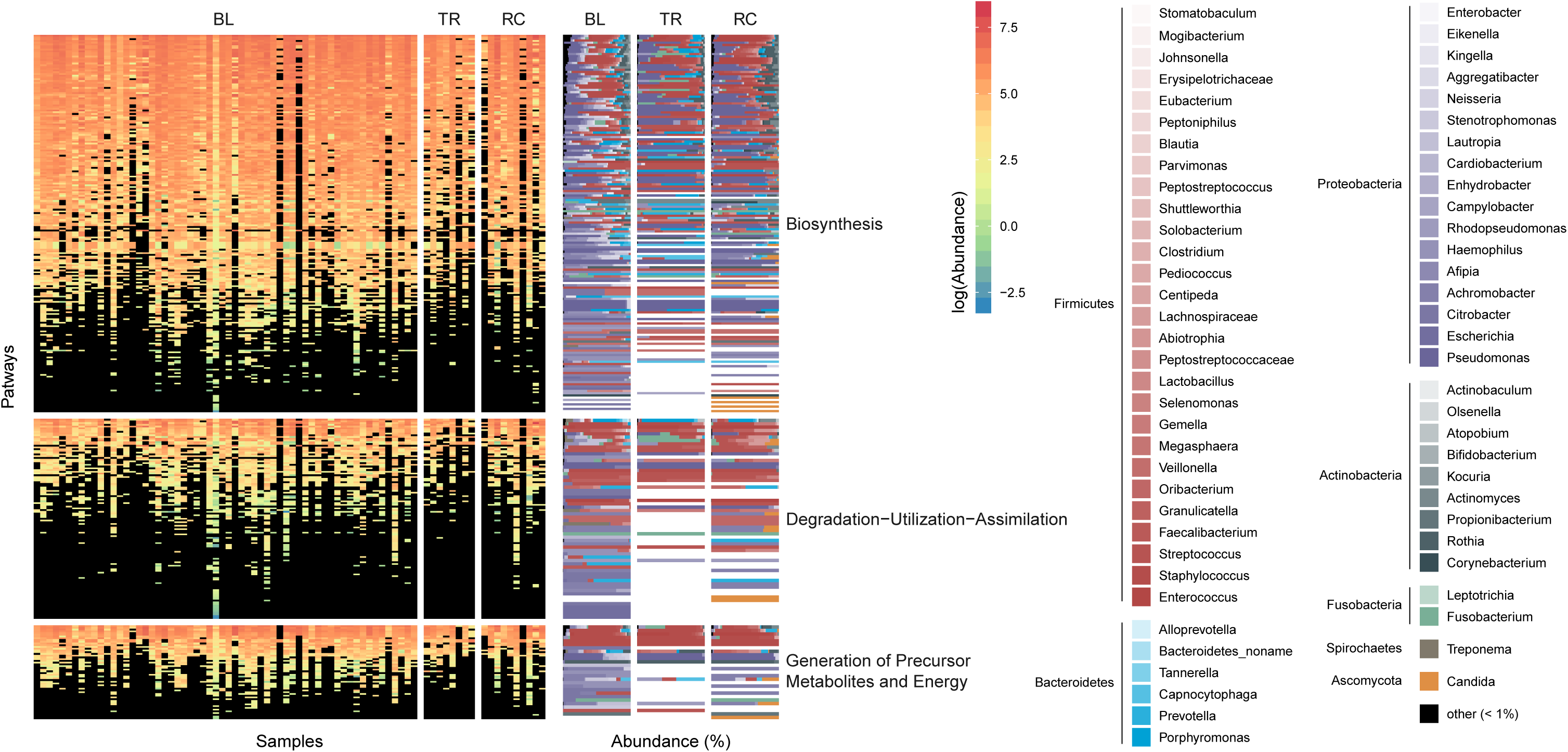
Pathway distribution according to exacerbation events. The pathway distribution was reported for each sample (columns) and for each pathway detected (rows). Colors from dark blue to red were used to report “copies per million” (CPM) values as obtained from HUMAnN2 with black reporting a CPM value of zero. On the left, the percentage of taxa in which each pathway was detected was reported using different colors. The main colors correspond to the Phylum whereas the different shades correspond to the genus detected (if available). BL, baseline; TR, treatment; RC, recovery.

**FIGURE 6.**
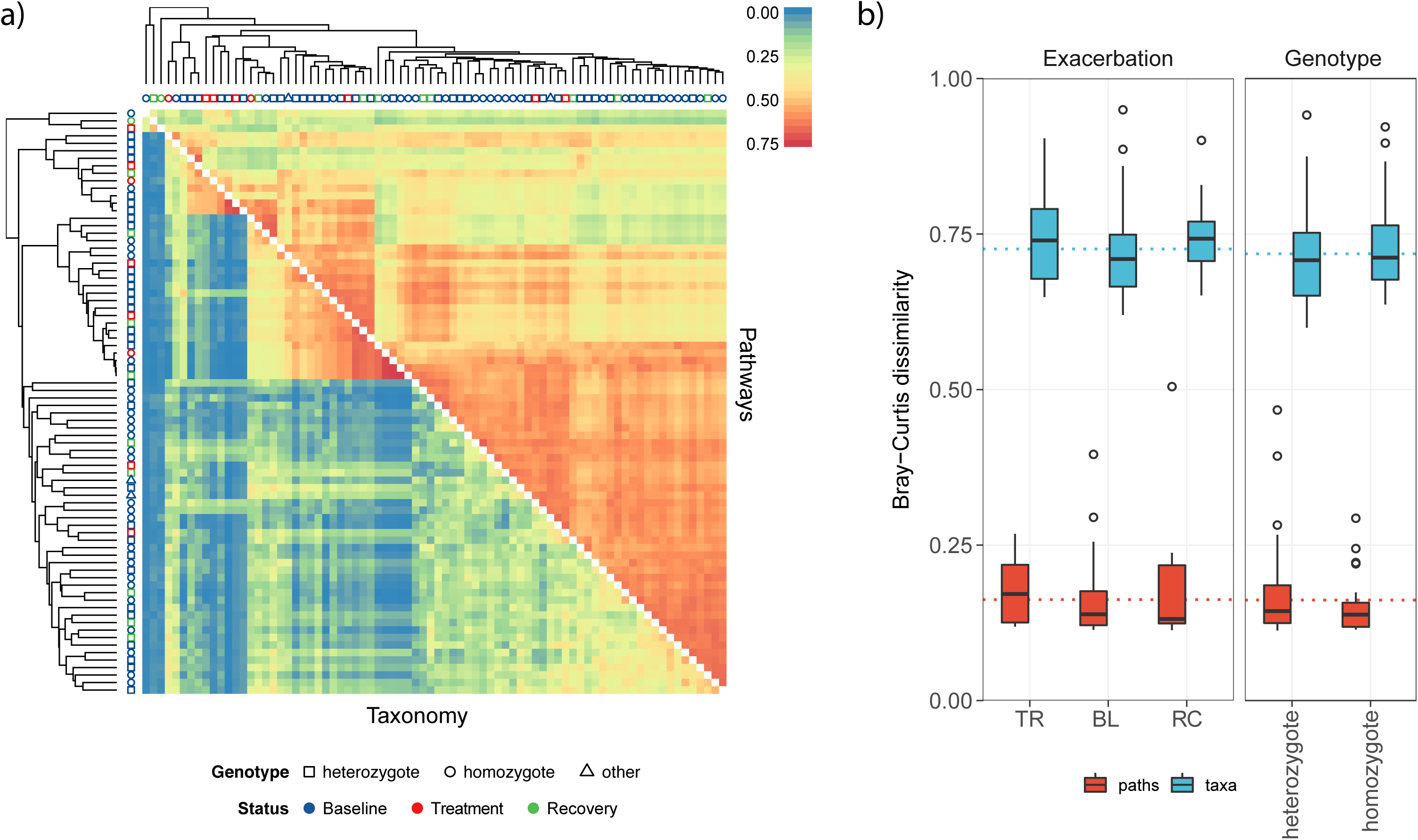
Beta diversity analysis on both taxonomic and functional distribution. a) Hierarchical clustering based on UPGMA method. Clustering was performed on both pathway distribution (the upped triangle) and taxonomic composition of samples (lower triangle). The Bray-Curtis distance was used to compute distances between samples, but it was transformed into similarity value by subtracting 1 before plotting. Thus, red colors represent high similarity values whereas blue colors represent low similarity values. The shape of the points on each tip of trees refers to the hospital whereas the colors refer to the exacerbation events. b) Results of Tukey’s post hoc test on beta diversity values across patient genotypes and exacerbation events. Contrasts were computed even to test differences between taxonomic distribution and pathways with taxa reporting higher level of beta diversity. Homozygote and heterozygote refer to ΔF508 mutation of CFTR gene. BL, baseline; TR, treatment; RC, recovery. Boxplot were computed as described in Figure 4 legend.

### Antibiotic resistance genes through exacerbation events and treatments

Similar to the pathway analysis reported above, antibiotic resistance genes (ARG) were inspected in relation to treatment events. Only six genes were found to be affected by an exacerbation condition, all regarding samples from patients heterozygous for ΔF508 whereas, as found for metabolic pathways, no gene was significantly impacted in terms of abundance by antibiotic treatment when considering all samples at once (supplementary materials Fig. S4 and Table S8). A similar approach was used to inspect the effect of antibiotic treatment on ARG distribution. ARG were inspected in relation to the antibiotic treatments reported in supplementary materials Table S1. The class of each antibiotic was correlated to the presence (and the abundance) of genes that may, in principle, confer resistance to antibiotics from the corresponding class. Differential abundance analyses were performed for each classes of antibiotics that was used in this study and results obtained were reported in supplementary materials Fig. S5 and Table S9. Only 11 genes were found to be affected by antibiotic intake in different ways. Indeed, 8 out of 11 reported a reduction of abundance during the treatment whereas the remaining 3 reported an increased abundance in respect with antibiotic intake. Results obtained confirmed the high resilience of the gene composition of CF lung microbiome. The relationship between presence of ARGs with antibiotic treatment was also explored. Results showed a large group of ARGs present in most of the samples and that the antibiotic treatment used in each sample was mirrored by the representation of the ARG classes (Supplementary materials Fig. S6 and Fig. S7).

## Discussion

Longitudinal studies provide important information on the stability and dynamics of microbial ecosystems (23). Here, we investigated the temporal dynamics of the CF sputum microbiome using shotgun metagenomics, including both periods of stability and respiratory exacerbations. The sputum microbiomes of CF patients were highly patient-specific and were substantially impacted both by the host and its lifestyle, suggesting the host has one of the most important determinants of sputum microbiome composition. Indeed, there was less variation within the same individual at different time points than between different individuals at the same time point, proving some degree of temporal stability of an individual’s sputum microbiome, as indicated by the lack of a time effect on the taxonomic distribution of microbiomes. The predominant taxa detected in sputa of CF patients exhibited extraordinary resilience, as demonstrated by the presence of the same strains of several species during the entire study period. While similar conclusions have been drawn from previous studies using both culture and 16S rRNA gene profiling, these studies failed to report a comprehensive, taxonomy-wide view of strain dynamics due to the limitations of these approaches (4, 6, 24). Carmody and colleagues showed a relatively stable sputum community that was often altered during period of exacerbation even in the absence of viral infection or antibiotic only in a small group of patients (5). A similar result was shown in the work from Fodor and colleagues (4) where, though occasional short-term compositional changes in the airway microbiota were found, the main taxonomic signatures of CF disease were highly stable. Even in other pulmonary diseases, such as non-cystic fibrosis bronchiectasis, respiratory sample bacterial communities showed a conserved structure for long period of time, as showed in the work by Cox and colleagues where patients were followed for a six-month period (24). In our study a notable exception was found for *Rothia mucilaginosa*. In fact, in contrast with other studies where this species was rarely identified (25–27), in our samples it was detected in high relative abundance. This finding may suggest a potential involvement of *R. mucilaginosa* in CF microbiome dynamics and pathogenicity, which deserves further attention.

Antibiotic exposure did not result in durable, persistent changes in sputum microbiota; the main taxa linked to CF infection were still present even after aggressive antibiotic treatment. From a taxonomic perspective, samples coming from the same patient clustered together, highlighting the role of the host in bacterial strain selection during the baseline but even during (and after) exacerbation events. Strain selection can be indeed influenced by a number of environmental factors (such as pH level and/or availability of nutrients) that are specific of the lung of a given host and cannot be determined *a-priori*. Despite this patient-specific colonization, sputum taxonomic composition differed significantly from one subject to another even when sampled at the same time. Conversely, microbial functional genetic pathways were more homogeneous across patients. This high conservation could indicate common features of the CF lung environment itself, such as mucus composition, nutrient availability, and oxygen levels, which can be broadly similar across patients with a similar clinical status. Such interpretation is consistent with the concept that the function of a biotic community is more conserved than the presence of single members due to functional redundancy of different microbial taxa (28). In fact, though the overall sputum microbiome in our study population included large set of microorganisms, the main functions detected are similar across all patients. From this point of view the airway microbiome can be considered as performing a similar “ecosystem service”, irrespective of the taxonomy present as pointed out by various authors in other environments (28, 29). The finding that CFTR genotypes relate with different representation in some pathways, may suggest that the airways microbiome is influenced by the type of CFTR alteration. However, this hypothesis deserves further attention to clarify a putative role of microbial pathways with respect to CFTR genotype and viceversa. Despite a clear effect of antibiotic treatment during (and after) exacerbation periods, the community structure is always recovered with the main pathogenic taxa emerging again. This effect is confirmed by the correlation of ARG distribution and antibiotic intake. Patients subjected to a given antibiotic treatment did not seem to select bacteria resistant to the antibiotic used but the detection of a particular mechanism seems to be distributed in almost all patients regardless of the treatment. An evidence of functional stability of the lung microbiota was previously reported in other works not concerning CF disease (30, 31). Both works focused their attention on the gut microbiome of obese and healthy individuals (human and mouse) reporting a considerable metabolic redundancy. This high degree of redundancy in the gut microbiome supported a more ecological view where subjects can be considered to some extent as different ecological niches, inhabited by unique collections of microbial phylotypes, but sharing the same set of genes. This concept can be applied to our cohort of patients, whose lung microbiome was taxonomically variable over time and among individuals (tough all chronically infected by the same species, *P. aeruginosa*), but where it was possible to identify a core set of metabolic-related gene features. This functional conservation may thus be needed by the whole community and patients can be seen as multiple micro-environments inhabited by a peculiar set of strains, which share the same functions. Investigations on the actual functionality (e.g. by metatranscriptomics) of the identified core-set of genes could provide clues on genetic function of the microbiome to be targeted in future therapeutic treatments (10). Additionally, the observed relations of pathway representation with CFTR genotype, though needing to be validated in larger studies, could offer possible opportunities for treating patients by targeting some CFTR genotype-related microbial metabolism. In conclusion, the temporal dynamics of the sputum microbiome in the largest cohort of patients with CF revealed analysed so far, showed patient-specific signatures of the airway microbiome at strain-level, lack of variation in the microbiome across pulmonary exacerbations, and a core set of antibiotic resistance genes that did not vary by antibiotic intake.

## Materials and Methods

### Ethics Statement

The study was approved by the Ethics Committees of Children’s Hospital and Research Institute Bambino Gesù (Rome, Italy), Cystic Fibrosis Center, Anna Meyer Children’s University Hospital (Florence, Italy) and G. Gaslini Institute (University of Genoa, Genoa, Italy) [Prot. N. 681 CM of November 2, 2012; Prot. N. 85 of February 27, 2014; Prot. N. FCC 2012 Partner 4-IGG of September 18, 2012]. All participants provided written informed consent before the enrollment in the study. All sputum specimens were produced voluntarily. All procedures were performed in agreement with the “Guidelines of the European Convention on Human Rights and Biomedicine for Research in Children” and the Ethics Committee of the three CF Centers involved. All measures were obtained and processed ensuring patient data protection and confidentiality.

### Characteristics of enrolled patients

Twenty-two adolescents and adults with moderate-severe lung disease and carrying the ΔF508 mutation were enrolled in the study between October 2014 and March 2015 (Table 1). The inclusion criteria are described in detail in the supplementary methods. Clinical status at the time of collection was designated as *baseline* (BL), when clinically stable and at their clinical and physiological baseline, *on treatment* (TR), at exacerbation-associated antibiotic treatments, and *at recovery* (RC), upon completion of antibiotic treatment. Subjects were treated according to current standards of care with periodical microbiological controls (32) with at least four microbiological controls per year (2). At each visit, clinical data collection and microbiological status (colonizing pathogens with available cultivation protocols) were performed according to the European CF Society standards of care (33). Forced expiratory volume in 1 second as a percentage of predicted (%FEV_1_) is a key outcome of monitoring lung function in CF (34). FEV_1_ values were measured according to the American Thoracic Society and European Respiratory Society standards (32). CFTR genotype, sex, age, and antibiotic treatment for each patient were reported in (Table 1 and supplementary materials Table S1). During serial sampling, data (antibiotic usage and spirometry) were collected. A total of 79 sputum sample were collected and DNA extraction were performed as reported in supplementary methods.

### Bioinformatic analyses

Sequence quality was ensured by trimming reads using StreamingTrim 1.0 (35), with a quality cutoff of 20. Bowtie2 (36) was used to screen out human-derived sequences from metagenomic data with the latest version of the human genome available in the NCBI database (GRCh38) as reference. Sequences displaying a concordant alignment (mate pair that aligns with the expected relative mate orientation and with the expected range of distances between mates) against the human genomes were then removed from all subsequent analyses. Metabolic and regulatory patterns were estimated using HUMAnN2 (37) and considering only those pathways with a coverage value ≥ 80%, whereas ≥ the taxonomic microbial community composition was assessed using MetaPhlAn2 (38). Reads were assembled into contigs using the metaSPAdes microbial assembler (39) with automatic k-mer length selection. To establish an airway microbiome gene catalog (7) we first removed contigs smaller than 500bp and then used prodigal in Anonymous mode (40), as suggested by the author of the tool, to predict open reading frames (ORFs). Translated protein sequences obtained from assembled contigs were classified using eggNOG mapper against the bactNOG database (41). Each protein was classified according to its best hit with an e-value lower than 0.001 as suggested in (42). The CARD database (43) was used in combination to the Resistance Gene Identifier (RGI, version 4.0.3) to inspect the distribution of antibiotic resistance gene (AR genes). Genes predicted within each metagenome were quantified using the number of reads that mapped against metagenomic contigs obtained for each sample. Reads were mapped back to contigs using Bowtie2 (36) and the number of reads mapping each ORF was obtained with the bedtools command “multicov” (version 2.26.0). To quantify gene content across different samples, genes were collapsed using the bestOG given by eggNOG mapper by summing together the number of reads that mapped genes with the same annotation. The same approach was used to quantify AR genes predicted with RGI but this time the unique identifier provided by CARD was used to collapse counts.

Strain characterization was performed using StrainPhlAn (20). Sequence variants for each organism detected were assessed against the MetaPhlAn2 (38) marker genes and a tree has been generated including all samples in which the organism was found at least in one time point. Since all organisms detected had at least one reference genome available in the RefSeq database, the most recent version of their genome was downloaded and added to the tree.

## Acknowledgments

We greatly acknowledge Professor Luke Hoffman (University of Washington, School of Medicine, Seattle, WA, USA) for critically reading and reviewing the earlier drafts of the manuscript. We also thank the Italian Cystic Fibrosis Research Foundation (FCC) (https://www.fibrosicisticaricerca.it) for financial support and administrative tasks, and the medical student Marta Maggisano (Sant’Andrea Hospital, Sapienza University of Rome, Rome, Italy) for the stimulating discussion. We would like to express their gratitude to Ricciotti Gabriella (Children’s Hospital and Research Institute Bambino Gesù, Rome), Dr. Tuccio Vanessa (Children’s Hospital and Research Institute Bambino Gesù, Rome) and Dr. Campana Silvia (Cystic Fibrosis Center, Anna Meyer Children’s University Hospital, Department of Pediatrics Medicine, Florence) for their technical support.

## Availability of data and material

All data generated or analysed during this study are included in this article. Raw sequence data reported in this study have been deposited in the NCBI “Sequence Read Archive” (SRA) under the project accession PRJNA516870.

## Authors’ contributions

Conceived and designed the experiments: AB VL GT EVF AM. Performed the experiments: GB FDC. Analyzed the data: GB AM AB. Contributed reagents/materials/analysis tools: DD FA PM RS AN. Wrote the paper: GB AM AB. Provided comments and recommendations that improved the manuscript: NS GT VL. Supervised research: AB VL GT EVF AM.

## Supplementary materials

Supplementary methods

### SUPPLEMENTARY TABLES

**TABLE S1** CFTR genotype, sex, age, and antibiotic treatment for each patient

**TABLE S2** Number of reads for each sample

**TABLE S3** Summary of all species detected with a mean abundance higher than 0.2%

**TABLE S4** Analysis of variance on alpha diversity indices

**TABLE S5** Tukey post hoc test on alpha diversity indices

**TABLE S6** Tukey post hoc test on Sorensen similarity index

**TABLE S7** Metabolic pathways differentially distributed across clinical statuses

**TABLE S8** Antibiotic resistance genes differentially distributed across clinical statuses

**TABLE S9** Antibiotic resistance genes differentially distributed depending on drug intake

### SUPPLEMENTARY FIGURES

**FIGURE S1** Strain-level phylogenetic trees of all detected microbes in the study

**FIGURE S2** Effect of genotypes and samples on the bacterial diversity of lung microbiome

**FIGURE S3** Differential abundant pathways

**FIGURE S4** Differential abundant antibiotic resistance genes

**FIGURE S5** Effect of the antibiotic intake on the distribution of antibiotic resistance genes

**FIGURE S6** Antibiotic resistance genes map

**FIGURE S7** Antibiotic resistance map of each sample included in the study

