## Supplementary materials for "Taxonomic variability over functional stability in the microbiome of Cystic Fibrosis patients chronically infected by *Pseudomonas aeruginosa*"

**Respiratory microbiome dynamics in CF patients with chronic *P. aeruginosa* infection**

50139, Italy

^3^Centre for Integrative Biology, University of Trento, Trento, 38122, Italy

^4^Children’s Hospital and Research Institute Bambino Gesù, Rome, 00165, Italy

^5^Cystic Fibrosis Center, IRCCS G. Gaslini Institute, Department of Pediatrics, Genoa, 16147, Italy

^6^Department for Sustainability, Italian National Agency for New Technologies, Energy and Sustainable Economic Development, ENEA, 00123 Rome, Italy

* Corresponding author

Annamaria Bevivino, Department for Sustainability, ENEA, Rome, Italy.

**Supplementary Methods**

### Inclusion criteria

The study subjects were selected based on eligibility criteria that included all of the following: (i) a diagnosis of CF, i.e., a sweat test showing sweat Cl > 60 mmol/l and two known CFTR mutations causing the disease with pancreatic insufficiency (elastase< 5 μg/g/feces) [1], (ii) aged more than six years, i.e., between 11 and 55 years, (iii) chronically infected with *Pseudomonas aeruginosa* according to the Leeds criteria [2] and iv) decline in %FEV_1_ in the previous three years before enrollment by measuring the difference between the best %FEV_1_ registered within the previous year and the best %FEV_1_ registered two-years before specimen collection, following the criteria previously reported [3]. Patients were excluded if they were chronically infected with *Burkholderia cepacia* complex. The cohort was enrolled in three Italian Hospital, namely: Bambino Gesù Children's Hospital (Rome, Italy), G. Gaslini Children's Hospital (Genoa, Italy) and Meyer Children's Hospital (Florence, Italy).

### Sample collection, processing, DNA extraction and sequencing

Sputum samples were obtained by spontaneous expectoration at baseline, exacerbation-associated antibiotic treatments and recovery status. Sampled were processed according to standard methods as previously described [4, 5]. Bacterial respiratory pathogens were identified using the conventional techniques reported in the Guidelines, as previously described [4, 6]. The number of samples, microbiological status at sampling and samplings following exacerbation events are reported in Table 1. Sputum samples were washed in 5 ml PBS and then centrifuged (3,800 g) for 15 minutes. Resulting pellets were resuspended in 5-10 ml DNAse buffer (10 mM Tris-HCl pH 7.5; 2.5 mM MgCl2; 0.5 mM CaCl2, pH 6.5) with 7.5 ul of DNAse I (2000 Units/ml) per 1 ml of sample (15U/ml final), incubated for 2 hours at 37C, and washed twice by pelleting at 3,800 g for 15 minutes and resuspending in 10 ml SE buffer (75 mM NaCl, 25 mM EDTA, pH 7.5). Pellets were then resuspended in 0.5 ml lysis buffer (20 mM Tris-HCl pH 8.0; 2 mM EDTA pH 8.0; 1% (v/v) Triton; 20 mg/ml Lysozyme final concentration), incubated for 30 minutes at 37C before extracting DNA with the MoBio Powersoil DNA extraction kit as per manufacturer's instructions. Libraries were prepared with Nextera XT kit (Illumina) Sequencing was performed on an Illumina HiSeq2500 apparatus (Illumina). Raw sequence data reported in this study have been deposited in the NCBI “Sequence Read Archive” (SRA) under the project accession PRJNA516870.

### Taxonomic classification of metagenomic contigs

Assembled contigs were taxonomically classified using BLAST. First, all genomes available for each species detected with MetaPhlAn2 were downloaded from NCBI and used to build a database for each sample. All genomes reporting an identity higher than 90% and a coverage higher than 80% were collected and used for taxonomic classification. Contigs reporting hits with genomes coming from a single species were assigned to that species whereas contigs reporting hits from multiple species were flagged as unknown.

### Statistical analyses

Statistical analyses were performed in R [7] version 3.4.4. The taxonomical and functional composition on lung microbiome was explored using permutational multivariate analysis of variance (PERMANOVA with 1000 permutations), ‘adonis2’ function of vegan package version 2.5-2; whereas differences in bacterial diversity were tested using analysis of covariance (ANCOVA), ‘aov’ function. The model fitted for both analyses was:

X ~ Status + Genotype + Subject + FEV_1_ + days

where, Exacerbation is the exacerbation event, Genotype is the CFTR genotype, Subject is the patient, FEV_1_ was the forced expiatory volume in 1 second, and days, was the number of days from the enrollment in the study. For the ANCOVA analyses Tukey's post hoc tests were performed to test for mean differences within each factor used to build the full model (excluding FEV_1_ value and days since they were not categorical variable). Ordination analyses were conducted on both taxa and pathways using the function ‘ordinate’ of the phyloseq package (version 1.23.1) with principle coordinate decomposition method (PCoA) and the Bray-Curtis dissimilarity index. The same index was used to inspect the distribution of samples and compare beta diversity level in bot taxonomic composition and pathways.

To test for differentially distributed pathways and taxa across exacerbation events and genotypes we used a moderated t-test as implemented in the limma package [8], version 3.34.9. Data obtained with MetaPhlAn2 (taxonomic composition) and HUMAnN2 (pathway composition) were fitted into limma’s model using subjects as blocking variable. Since both software quantify biological units using relative counts (HUMAnN2 uses “copies per million” and MetaPhlAn2 uses percentages) we transformed this data into logarithmic values using the formula: log_2_(x + 0.1), where x are the relative counts. Obtained p-values were corrected using the Benjamini-Hochberg correction method. A similar approach has been used for antibiotic genes detect along assembled contigs. Here the number of reads that mapped onto each gene was used to estimate differentially abundant gene. Since the number of reads for each sample was variable (the ratio of the largest library size to the smallest was more than 10-fold) we used limma’s voom method [9] to fit our model, as suggested by the author of limma.

**Supplementary Tables**

**TABLE S2**

Number of reads for each sample

| ID | Time point | # reads | # pairs | # unpaired |
| --- | --- | --- | --- | --- |
| B01 | 0 | 5503844 | 2325382 | 853080 |
| B01 | 1 | 868905 | 406838 | 55229 |
| B01 | 2 | 504486 | 214496 | 75494 |
| B01 | 3 | 1123511 | 530841 | 61829 |
| B01 | 1 | 1288912 | 623800 | 41312 |
| B02 | 0 | 1502243 | 623532 | 255179 |
| B02 | 2 | 8991858 | 4451619 | 88620 |
| B02 | 3 | 5143954 | 2488108 | 167738 |
| B03 | 0 | 658063 | 190398 | 277267 |
| B03 | 1 | 2115889 | 1025708 | 64473 |
| B03 | 2 | 10714542 | 5261449 | 191644 |
| B03 | 3 | 7484908 | 3638635 | 207638 |
| B06 | 0 | 5069581 | 2234180 | 601221 |
| B06 | 1 | 2070381 | 996544 | 77293 |
| B06 | 2 | 13180766 | 6517309 | 146148 |
| B06 | 3 | 24111405 | 11622361 | 866683 |
| G10 | 0 | 10932913 | 4877651 | 1177611 |
| G10 | 1 | 14514555 | 7134118 | 246319 |
| G10 | 2 | 3628618 | 1751868 | 124882 |
| G10 | 3 | 5452496 | 2625377 | 201742 |
| G24 | 0 | 785844 | 320254 | 145336 |
| G24 | 1 | 995004 | 435853 | 123298 |
| G24 | 2 | 3536639 | 1728879 | 78881 |
| G28 | 0 | 5760251 | 2518915 | 722421 |
| G28 | 1 | 661864 | 321914 | 18036 |
| G30 | 0 | 1836789 | 678840 | 479109 |
| G31 | 0 | 242296 | 102687 | 36922 |
| G31 | 1 | 1777716 | 859179 | 59358 |
| G34 | 1 | 605036 | 281046 | 42944 |
| M05 | 0 | 12125170 | 5264493 | 1596184 |
| M05 | 1 | 1124272 | 496383 | 131506 |
| M05 | 2 | 1470732 | 701131 | 68470 |
| M05 | 3 | 1153167 | 533168 | 86831 |
| M19 | 0 | 6259797 | 2704837 | 850123 |
| M19 | 1 | 17369379 | 8572310 | 224759 |
| M19 | 2 | 9391369 | 4639094 | 113181 |
| M19 | 3 | 9318334 | 4611450 | 95434 |
| M21 | 0 | 3128755 | 1369530 | 389695 |
| M21 | 1 | 224963 | 74970 | 75023 |
| M21 | 2 | 1335393 | 628615 | 78163 |
| M21 | 1 | 216831 | 58002 | 100827 |
| M22 | 0 | 3088535 | 1311593 | 465349 |
| M22 | 1 | 806490 | 381277 | 43936 |
| M22 | 2 | 1496116 | 721477 | 53162 |
| M22 | 3 | 819739 | 321497 | 176745 |
| M22 | 1 | 831267 | 394417 | 42433 |
| M23 | 0 | 964452 | 413639 | 137174 |
| M23 | 1 | 2031131 | 986181 | 58769 |
| M23 | 2 | 991655 | 466705 | 58245 |
| M23 | 1 | 970320 | 456575 | 57170 |
| M24 | 0 | 6030041 | 2542675 | 944691 |
| M24 | 1 | 922 | 446 | 30 |
| M24 | 2 | 31295587 | 15469995 | 355597 |
| M24 | 3 | 16800705 | 8254483 | 291739 |
| M25 | 0 | 18762529 | 8411793 | 1938943 |
| M25 | 1 | 5227170 | 2569092 | 88986 |
| M25 | 2 | 535041 | 181216 | 172609 |
| M25 | 3 | 7454111 | 3658164 | 137783 |
| M26 | 0 | 2258264 | 979675 | 298914 |
| M26 | 1 | 8457325 | 4176615 | 104095 |
| M26 | 2 | 5689324 | 2797112 | 95100 |
| M26 | 3 | 2919836 | 1413367 | 93102 |
| M26 | 1 | 33719580 | 16657112 | 405356 |
| M28 | 0 | 3458580 | 1524503 | 409574 |
| M28 | 1 | 5827736 | 2853068 | 121600 |
| M28 | 2 | 3070091 | 1440938 | 188215 |
| M28 | 3 | 1029948 | 475535 | 78878 |
| M29 | 0 | 1500423 | 599852 | 300719 |
| M29 | 1 | 936022 | 411010 | 114002 |
| M29 | 2 | 2076152 | 995563 | 85026 |
| M29 | 3 | 521984 | 183202 | 155580 |
| M31 | 0 | 6846501 | 2877219 | 1092063 |
| M31 | 1 | 1238914 | 552508 | 133898 |
| M31 | 2 | 1391941 | 664028 | 63885 |
| M33 | 0 | 1188194 | 434000 | 320194 |
| M33 | 1 | 3693755 | 1810992 | 71771 |
| M33 | 2 | 3108154 | 1492259 | 123636 |
| M33 | 3 | 2562448 | 1246795 | 68858 |
| M33 | 1 | 5604678 | 2758664 | 87350 |

ID, sample’s ID; Time point, time point at which sample was collected; # reads, total number of reads after preprocessing; # pairs, number of paired reads; # unpaired, number of unpaired reads.

**TABLE S3**

Summary of all species detected with a mean abundance higher than 0.2%

| Kingdom | Phylum | Class | Order | Family | Genus | Species | (Mean abundance ± standard error) % |
| --- | --- | --- | --- | --- | --- | --- | --- |
| Viruses | - | - | Caudovirales | Siphoviridae | Lambdalikevirus | Enterobacteria_phage_lambda | 1.06 ± 0.508 |
|  |  |  |  | Podoviridae | Podoviridae_noname | Streptococcus_phage_Cp_1 | 0.30 ± 0.213 |
| Bacteria | Proteobacteria | Gammaproteobacteria | Pseudomonadales | Pseudomonadaceae | Pseudomonas | Pseudomonas_unclassified | 1.18 ± 0.494 |
|  |  |  |  |  |  | **Pseudomonas_aeruginosa** | 25.44 ± 3.657 |
|  |  |  | Pasteurellales | Pasteurellaceae | Haemophilus | Haemophilus_parainfluenzae | 0.95 ± 0.268 |
|  |  |  |  |  |  | Haemophilus_influenzae | 0.41 ± 0.331 |
|  |  | Betaproteobacteria | Neisseriales | Neisseriaceae | Neisseria | Neisseria_unclassified | 0.69 ± 0.308 |
|  |  |  |  |  |  | Neisseria_flavescens | 0.52 ± 0.226 |
|  |  |  | Burkholderiales | Burkholderiaceae | Ralstonia | Ralstonia_unclassified | 0.55 ± 0.258 |
|  |  |  |  |  | Lautropia | Lautropia_mirabilis | 0.22 ± 0.167 |
|  | Fusobacteria | Fusobacteriia | Fusobacteriales | Fusobacteriaceae | Fusobacterium | Fusobacterium_nucleatum | 0.30 ± 0.069 |
|  | Firmicutes | Negativicutes | Selenomonadales | Veillonellaceae | Veillonella | **Veillonella_unclassified** | 2.77 ± 0.470 |
|  |  |  |  |  |  | Veillonella_parvula | 1.76 ± 0.434 |
|  |  |  |  |  |  | Veillonella_dispar | 0.34 ± 0.093 |
|  |  |  |  |  |  | Veillonella_atypica | 0.66 ± 0.155 |
|  |  | Erysipelotrichia | Erysipelotrichales | Erysipelotrichaceae | Solobacterium | Solobacterium_moorei | 0.41 ± 0.118 |
|  |  | Bacilli | Lactobacillales | Streptococcaceae | Streptococcus | Streptococcus_salivarius | 0.45 ± 0.188 |
|  |  |  |  |  |  | **Streptococcus_parasanguinis** | 7.02 ± 1.091 |
|  |  |  |  |  |  | Streptococcus_mitis_oralis_pneumoniae | 1.19 ± 0.217 |
|  |  |  |  |  |  | Streptococcus_infantis | 1.11 ± 0.266 |
|  |  |  |  |  |  | Streptococcus_australis | 0.65 ± 0.196 |
|  |  |  | Lactobacillales | Enterococcaceae | Enterococcus | **Enterococcus_faecalis** | 2.45 ± 1.421 |
|  |  |  |  | Carnobacteriaceae | Granulicatella | Granulicatella_unclassified | 1.87 ± 0.269 |
|  |  |  |  |  |  | Granulicatella_adiacens | 1.19 ± 0.157 |
|  |  |  |  | Aerococcaceae | Abiotrophia | Abiotrophia_defectiva | 0.40 ± 0.187 |
|  |  |  | Bacillales | Staphylococcaceae | Staphylococcus | **Staphylococcus_aureus** | 11.19 ± 2.833 |
|  |  |  |  | Bacillales_noname | Gemella | **Gemella_sanguinis** | 1.89 ± 0.386 |
|  |  |  |  |  |  | Gemella_morbillorum | 0.51 ± 0.203 |
|  |  |  |  |  |  | Gemella_haemolysans | 1.39 ± 0.373 |
|  | Bacteroidetes | Flavobacteriia | Flavobacteriales | Flavobacteriaceae | Capnocytophaga | Capnocytophaga_unclassified | 0.41 ± 0.097 |
|  |  |  |  |  |  | Capnocytophaga_sp_oral_taxon_329 | 0.34 ± 0.214 |
|  |  |  |  |  |  | Capnocytophaga_gingivalis | 0.37 ± 0.105 |
|  |  | Bacteroidia | Bacteroidales | Prevotellaceae | Prevotella | Prevotella_sp_C561 | 0.23 ± 0.153 |
|  |  |  |  |  |  | Prevotella_pleuritidis | 0.62 ± 0.336 |
|  |  |  |  |  |  | Prevotella_pallens | 0.21 ± 0.056 |
|  |  |  |  |  |  | Prevotella_nanceiensis | 0.51 ± 0.207 |
|  |  |  |  |  |  | **Prevotella_melaninogenica** | 2.89 ± 0.733 |
|  |  |  |  |  |  | Prevotella_histicola | 0.77 ± 0.397 |
|  |  |  |  | Porphyromonadaceae | Porphyromonas | **Porphyromonas_sp_oral_taxon_279** | 4.03 ± 0.947 |
|  | Actinobacteria | Actinobacteria | Coriobacteriales | Coriobacteriaceae | Atopobium | Atopobium_parvulum | 0.21 ± 0.047 |
|  |  |  | Bifidobacteriales | Bifidobacteriaceae | Bifidobacterium | Bifidobacterium_longum | 0.38 ± 0.351 |
|  |  |  | Actinomycetales | Micrococcaceae | Rothia | **Rothia_mucilaginosa** | 9.07 ± 1.378 |
|  |  |  |  |  |  | **Rothia_dentocariosa** | 3.00 ± 0.568 |
|  |  |  |  |  |  | Rothia_aeria | 0.54 ± 0.246 |
|  |  |  |  | Actinomycetaceae | Actinomyces | Actinomyces_graevenitzii | 1.53 ± 0.478 |

Top 10 species detected were reported in bold.

**TABLE S4**

Analysis of variance on alpha diversity indices

|  | Df | Sum Sq | Mean Sq | F value | Pr(>F) |
| --- | --- | --- | --- | --- | --- |
| Taxonomic profiling | | | | | |
| *Shannon index* |  |  |  |  |  |
| Status | **2** | **3.57** | **1.78** | **3.73** | **0.0310** |
| Genotype | **1** | **2.34** | **2.34** | **4.89** | **0.0317** |
| Sample | **18** | **27.48** | **1.53** | **3.20** | **0.0006** |
| FEV_1_ value | 1 | 1.01 | 1.01 | 2.12 | 0.1521 |
| Days | 1 | 0.06 | 0.06 | 0.13 | 0.7208 |
| Status:Genotype | 1 | 1.69 | 1.69 | 3.55 | 0.0656 |
| Residuals | 49 | 23.41 | 0.48 |  |  |
| *Inverse Simpson index* |  |  |  |  |  |
| Status | **2** | **54.14** | **27.07** | **3.27** | **0.0466** |
| Genotype | **1** | **64.24** | **64.24** | **7.75** | **0.0076** |
| Sample | **18** | **554.16** | **30.79** | **3.71** | **0.0001** |
| FEV_1_ value | 1 | 20.81 | 20.81 | 2.51 | 0.1195 |
| Days | 1 | 0.17 | 0.17 | 0.02 | 0.8878 |
| Status:Genotype | 1 | 7.56 | 7.56 | 0.91 | 0.3442 |
| Residuals | 49 | 406.11 | 8.29 |  |  |
| Pathway distribution | | | | | |
| *Shannon index* |  |  |  |  |  |
| Status | 2 | 0.45 | 0.22 | 3.12 | 0.0533 |
| Genotype | 1 | 0.11 | 0.11 | 1.55 | 0.2189 |
| Sample | **18** | **3.90** | **0.22** | **3.03** | **0.0011** |
| FEV_1_ value | 1 | 0.19 | 0.19 | 2.62 | 0.1119 |
| Days | 1 | 0.08 | 0.08 | 1.06 | 0.3086 |
| Status:Genotype | 1 | 0.27 | 0.27 | 3.75 | 0.0587 |
| Residuals | 49 | 3.50 | 0.07 |  |  |
| *Inverse Simpson index* |  |  |  |  |  |
| Status | **2** | **4056.46** | **2028.23** | **3.57** | **0.0358** |
| Genotype | 1 | 269.70 | 269.70 | 0.47 | 0.4942 |
| Sample | **18** | **32244.56** | **1791.36** | **3.15** | **0.0007** |
| FEV_1_ value | 1 | 2010.30 | 2010.30 | 3.54 | 0.0660 |
| Days | 1 | 225.89 | 225.89 | 0.40 | 0.5314 |
| Status:Genotype | 1 | 2248.92 | 2248.92 | 3.96 | 0.0523 |
| Residuals | 49 | 27852.86 | 568.43 |  |  |

Results of analysis of variance (ANOVA) on Shannon and Inverse Simpson indices were reported in table. Results obtained from distribution of taxa and pathways were reported. Factors with a p-value lower than 0.05 were reported in bold.

**TABLE S5**

Tukey post hoc test on alpha diversity indices

|  | Index | Contrast | Mean diff. | 2.5% (C.I.) | 97.5% (C.I.) | p-value |
| --- | --- | --- | --- | --- | --- | --- |
| Taxonomic profiling | | | | | | |
| Genotype |  |  |  |  |  |  |
|  | Shannon | HO – HE | 0.35 | 0.02 | 0.69 | 0.0405* |
|  | InvSimpson | HO – HE | 1.84 | 0.45 | 3.24 | 0.0108* |
| Status |  |  |  |  |  |  |
|  | Shannon | BL - TR | 0.67 | 0.04 | 1.30 | 0.0363* |
|  |  | RC - TR | 0.35 | -0.47 | 1.16 | 0.5631 |
|  |  | RC – BL | -0.32 | -0.92 | 0.28 | 0.4048 |
|  | InvSimpson | BL - TR | 2.52 | -0.11 | 5.15 | 0.0328* |
|  |  | RC - TR | 1.10 | -2.28 | 4.49 | 0.7113 |
|  |  | RC – BL | -1.41 | -3.91 | 1.08 | 0.3646 |
| Pathway distribution | | | | | | |
| Genotype |  |  |  |  |  |  |
|  | Shannon | HO – HE | 0.08 | -0.05 | 0.21 | 0.2416 |
|  | InvSimpson | HO – HE | 3.77 | -7.80 | 15.34 | 0.5151 |
| Status |  |  |  |  |  |  |
|  | Shannon | BL - TR | 0.22 | -0.02 | 0.47 | 0.0813 |
|  |  | RC - TR | 0.08 | -0.23 | 0.40 | 0.8017 |
|  |  | RC – BL | -0.14 | -0.37 | 0.09 | 0.3212 |
|  | InvSimpson | BL - TR | 21.45 | -0.30 | 43.21 | 0.0540 |
|  |  | RC - TR | 8.57 | -19.43 | 36.57 | 0.7409 |
|  |  | RC – BL | -12.88 | -33.54 | 7.79 | 0.2971 |

Factors were reported in bold whereas alpha diversity indices were reported once for all contrasts performed. Contrast were reported using the following abbreviations: HO, homozygote ΔF_508_; HE, heterozygote ΔF_508_; BL, baseline; TR, treatment (samples collected during the treatment of an exacerbation event); RC, recovery (the first sample collected after the end of an exacerbation event). The differences between the mean values of the given contrast were reported in the Mean diff. columns whereas the confidence interval was reported using the C.I. abbreviation. Contrasts reporting a p-value lower than 0.05 were marked with an asterisk.

**TABLE S6**

Tukey post hoc test on Sorensen similarity index

|  | Contrast | Mean diff. | 2.5% (C.I.) | 97.5% (C.I.) | p-value |
| --- | --- | --- | --- | --- | --- |
| Genotype |  |  |  |  |  |
|  | Taxa - Pathways | 0.56 | 0.53 | 0.58 | < 0.0001* |
|  | HO - HE_t_ | 0.00 | -0.03 | 0.03 | 0.9805 |
| Status |  |  |  |  |  |
|  | Taxa - Pathways | 0.56 | 0.54 | 0.58 | < 0.0001* |
|  | BL - TR | -0.03 | -0.07 | 0.01 | 0.2704 |
|  | RC - TR | 0.01 | -0.05 | 0.06 | 0.9662 |
|  | RC – BL | 0.03 | -0.01 | 0.07 | 0.1027 |

Grouping factors were reported in bold whereas contrasts were reported using the following abbreviations: HO, homozygote ΔF_508_; HE, heterozygote ΔF_508_; BL, baseline; TR, treatment (samples collected during the treatment of an exacerbation event); RC, recovery (the first sample collected after the end of an exacerbation event). The differences between the mean values of the given contrast were reported in the Mean diff. column whereas the confidence interval was reported using the C.I. abbreviation. Contrasts with a pvalue lower than 0.05 were marked with an asterisk.

**TABLE S7**

Metabolic pathways differentially distributed across exacerbation statuses

| Genotype | Contrast | MetaCyc name | Path description | logFC | AveExpr | t | P.Value | adj.P.Val |
| --- | --- | --- | --- | --- | --- | --- | --- | --- |
| Homozygote | TR Vs BL | PWY0-162 | superpathway of pyrimidine ribonucleotides de novo biosynthesis | -6.51 | 7.31 | -4.86 | 0.0000 | 0.0018 |
| Homozygote | TR Vs BL | PWY-6147 | 6-hydroxymethyl-dihydropterin diphosphate biosynthesis I | -6.71 | 7.17 | -4.45 | 0.0000 | 0.0043 |
| Homozygote | TR Vs BL | PWY-7539 | 6-hydroxymethyl-dihydropterin diphosphate biosynthesis III (Chlamydia) | -6.13 | 6.71 | -4.32 | 0.0000 | 0.0046 |
| Homozygote | TR Vs BL | PWY-7199 | pyrimidine deoxyribonucleosides salvage | -6.33 | 7.14 | -4.15 | 0.0001 | 0.0065 |
| Homozygote | TR Vs BL | PWY-6628 | superpathway of L-phenylalanine biosynthesis | -6.01 | 6.86 | -3.91 | 0.0002 | 0.0123 |
| Homozygote | TR Vs BL | DAPLYSINESYN-PWY | L-lysine biosynthesis I | -5.75 | 6.13 | -3.81 | 0.0003 | 0.0123 |
| Homozygote | TR Vs BL | PWY-6703 | preQ0 biosynthesis | -5.83 | 6.28 | -3.79 | 0.0003 | 0.0123 |
| Homozygote | TR Vs BL | PENTOSE-P-PWY | pentose phosphate pathway | -5.51 | 6.28 | -3.77 | 0.0003 | 0.0123 |
| Homozygote | TR Vs BL | PWY0-1319 | CDP-diacylglycerol biosynthesis II | -6.29 | 6.49 | -3.70 | 0.0004 | 0.0142 |
| Homozygote | TR Vs BL | PWY-7187 | pyrimidine deoxyribonucleotides de novo biosynthesis II | -5.78 | 6.48 | -3.54 | 0.0006 | 0.0191 |
| Homozygote | TR Vs BL | PWY-621 | sucrose degradation III (sucrose invertase) | -5.56 | 6.12 | -3.53 | 0.0007 | 0.0191 |
| Homozygote | TR Vs BL | DENOVOPURINE2-PWY | superpathway of purine nucleotides de novo biosynthesis II | -5.89 | 6.59 | -3.52 | 0.0007 | 0.0191 |
| Homozygote | TR Vs BL | PWY-5667 | CDP-diacylglycerol biosynthesis I | -5.91 | 6.24 | -3.46 | 0.0009 | 0.0216 |
| Homozygote | TR Vs BL | P4-PWY | superpathway of L-lysine, L-threonine and L-methionine biosynthesis I | -5.62 | 5.80 | -3.30 | 0.0014 | 0.0299 |
| Homozygote | TR Vs BL | PWY66-409 | superpathway of purine nucleotide salvage | -5.84 | 6.38 | -3.27 | 0.0016 | 0.0299 |
| Homozygote | TR Vs BL | MET-SAM-PWY | superpathway of S-adenosyl-L-methionine biosynthesis | -5.05 | 7.00 | -3.23 | 0.0018 | 0.0299 |
| Homozygote | TR Vs BL | RIBOSYN2-PWY | flavin biosynthesis I (bacteria and plants) | -5.03 | 5.73 | -3.15 | 0.0023 | 0.0319 |
| Homozygote | RC Vs BL | PWY-5188 | tetrapyrrole biosynthesis I (from glutamate) | -6.95 | 8.24 | -8.14 | 0.0000 | 0.0000 |
| Homozygote | RC Vs BL | HEMESYN2-PWY | heme b biosynthesis II (anaerobic) | -6.63 | 7.82 | -5.65 | 0.0000 | 0.0000 |
| Homozygote | RC Vs BL | ARGSYN-PWY | L-arginine biosynthesis I (via L-ornithine) | -6.33 | 7.59 | -5.42 | 0.0000 | 0.0001 |
| Homozygote | RC Vs BL | PWY-7400 | L-arginine biosynthesis IV (archaebacteria) | -6.21 | 7.54 | -5.12 | 0.0000 | 0.0002 |
| Homozygote | RC Vs BL | GLUTORN-PWY | L-ornithine biosynthesis I | -5.53 | 7.02 | -4.53 | 0.0000 | 0.0009 |
| Homozygote | RC Vs BL | PWY-5918 | superpathay of heme b biosynthesis from glutamate | -5.97 | 7.36 | -4.40 | 0.0000 | 0.0012 |
| Homozygote | RC Vs BL | RHAMCAT-PWY | L-rhamnose degradation I | 6.73 | 3.12 | 4.38 | 0.0000 | 0.0012 |
| Homozygote | RC Vs BL | HEME-BIOSYNTHESIS-II | heme b biosynthesis I (aerobic) | -5.53 | 7.09 | -4.16 | 0.0001 | 0.0023 |
| Homozygote | RC Vs BL | PWY-5189 | tetrapyrrole biosynthesis II (from glycine) | -5.05 | 7.33 | -3.95 | 0.0002 | 0.0044 |
| Homozygote | RC Vs BL | PWY0-162 | superpathway of pyrimidine ribonucleotides de novo biosynthesis | -6.47 | 7.31 | -3.45 | 0.0009 | 0.0161 |
| Homozygote | RC Vs BL | PWY-6147 | 6-hydroxymethyl-dihydropterin diphosphate biosynthesis I | -6.74 | 7.17 | -3.20 | 0.0019 | 0.0275 |
| Homozygote | RC Vs BL | PWY-6897 | thiamine salvage II | -6.28 | 6.54 | -3.10 | 0.0026 | 0.0355 |
| Homozygote | RC Vs BL | PWY-7539 | 6-hydroxymethyl-dihydropterin diphosphate biosynthesis III (Chlamydia) | -6.11 | 6.71 | -3.08 | 0.0028 | 0.0367 |
| Homozygote | RC Vs TR | PWY-5188 | tetrapyrrole biosynthesis I (from glutamate) | -6.84 | 8.24 | -7.17 | 0.0000 | 0.0000 |
| Homozygote | RC Vs TR | ARGSYN-PWY | L-arginine biosynthesis I (via L-ornithine) | -6.75 | 7.59 | -5.17 | 0.0000 | 0.0002 |
| Homozygote | RC Vs TR | RHAMCAT-PWY | L-rhamnose degradation I | 8.85 | 3.12 | 5.16 | 0.0000 | 0.0002 |
| Homozygote | RC Vs TR | PWY-7400 | L-arginine biosynthesis IV (archaebacteria) | -6.74 | 7.54 | -4.98 | 0.0000 | 0.0003 |
| Homozygote | RC Vs TR | HEMESYN2-PWY | heme b biosynthesis II (anaerobic) | -6.28 | 7.82 | -4.79 | 0.0000 | 0.0005 |
| Homozygote | RC Vs TR | GLUTORN-PWY | L-ornithine biosynthesis I | -5.94 | 7.02 | -4.36 | 0.0000 | 0.0020 |
| Homozygote | RC Vs TR | PWY-5189 | tetrapyrrole biosynthesis II (from glycine) | -5.47 | 7.33 | -3.83 | 0.0002 | 0.0090 |
| Homozygote | RC Vs TR | PWY-5918 | superpathay of heme b biosynthesis from glutamate | -5.74 | 7.36 | -3.79 | 0.0003 | 0.0094 |
| Homozygote | RC Vs TR | PWY-7199 | pyrimidine deoxyribonucleosides salvage | 8.52 | 7.14 | 3.57 | 0.0006 | 0.0162 |
| Homozygote | RC Vs TR | HEME-BIOSYNTHESIS-II | heme b biosynthesis I (aerobic) | -5.20 | 7.09 | -3.50 | 0.0008 | 0.0188 |

Results of limma analysis were reported in the table together with the genotype, the pathway name (according to MetaCyc), and the description of the pathway. Other columns are: logFC, the log2-transformed fold change value; AveExpr, the average log2-expression level for that pathway across all samples; t, the t-value according to limma’s moderated t-test; P.Value, the p-value; adj.P.Val, the adjusted p-value using the Benjamini–Hochberg correction. Contrast were reported using the following abbreviations: Homozygote, homozygote ΔF_508_; BL, baseline; TR, treatment (samples collected during the treatment of an exacerbation event); RC, recovery (the first sample collected after the end of an exacerbation event). Only contrasts with an adjusted p-value lower than 0.05 and an absolute log fold-change value higher than 5 were reported.

**TABLE S8**

Antibiotic resistance genes differentially distributed across exacerbation statuses

| Genotype | Contrast | Gene name | AR family | logFC | AveExpr | t | P.Value | adj.P.Val |
| --- | --- | --- | --- | --- | --- | --- | --- | --- |
| Heterozygote | RC Vs BL | PC1 beta-lactamase (blaZ) | blaZ beta-lactamase | 7.68 | 7.58 | 7.08 | 0.0000 | 0.0000 |
| Heterozygote | RC Vs BL | LRA-13 | class C LRA beta-lactamase; class D LRA beta-lactamase | 5.43 | 7.27 | 6.29 | 0.0000 | 0.0000 |
| Heterozygote | RC Vs TR | LRA-13 | class C LRA beta-lactamase; class D LRA beta-lactamase | 5.50 | 7.27 | 5.41 | 0.0000 | 0.0001 |
| Heterozygote | RC Vs TR | PC1 beta-lactamase (blaZ) | blaZ beta-lactamase | 7.91 | 7.58 | 5.26 | 0.0000 | 0.0001 |
| Heterozygote | RC Vs TR | sav1866 | ATP-binding cassette (ABC) antibiotic efflux pump | 6.50 | 8.78 | 3.89 | 0.0002 | 0.0050 |
| Heterozygote | RC Vs TR | tet(38) | major facilitator superfamily (MFS) antibiotic efflux pump | 6.95 | 9.00 | 3.60 | 0.0006 | 0.0118 |

Results of limma analysis were reported in the table together with the genotype, the antibiotic resistance gene name (according to the CARD database), and the antibiotic resistance family (AR family). Other columns are: logFC, the log2-transformed fold change value; AveExpr, the average log2-expression level for that pathway across all samples; t, the t-value according to limma’s moderated t-test; P.Value, the p-value; adj.P.Val, the adjusted pvalue using the Benjamini–Hochberg correction. Contrasts were reported using the following abbreviations: Homozygote, homozygote ΔF_508_; BL, baseline; TR, treatment (samples collected during the treatment of an exacerbation event); RC, recovery (the first sample collected after the end of an exacerbation event). Only contrasts with an adjusted p-value lower than 0.05 and an absolute log fold-change value higher than 5 were reported.

**TABLE S9**

Antibiotic resistance genes differentially distributed depending on drug intake

| Gene name | Gene Family | Resistance Mechanism | Drug Class | Antibiotic Class | logFC | AveExpr | t | P.Value | adj.P.Val |
| --- | --- | --- | --- | --- | --- | --- | --- | --- | --- |
| basS | pmr phosphoethanolamine transferase | antibiotic target alteration | peptide antibiotic | peptide antibiotic | -0.80 | 11.51 | -5.22 | <0.00001 | 0.0001 |
| FosA | fosfomycin thiol transferase | antibiotic inactivation | fosfomycin | peptide antibiotic | -1.10 | 10.08 | -3.56 | 0.0006 | 0.0199 |
| ArmR | resistance-nodulation-cell division (RND) antibiotic efflux pump | antibiotic efflux | aminocoumarin antibiotic; carbapenem; cephalosporin; cephamycin; diaminopyrimidine antibiotic; fluoroquinolone antibiotic; macrolide antibiotic; monobactam; penam; penem; peptide antibiotic; phenicol antibiotic; sulfonamide antibiotic; tetracycline antibiotic | peptide antibiotic | -1.56 | 9.24 | -3.42 | 0.0010 | 0.0213 |
| OXA-50 | OXA beta-lactamase | antibiotic inactivation | cephalosporin; penam | aminoglycoside antibiotic | -0.45 | 11.61 | -3.51 | 0.0008 | 0.0397 |
| Pseudomonas aeruginosa soxR | ATP-binding cassette (ABC) antibiotic efflux pump; major facilitator superfamily (MFS) antibiotic efflux pump; resistance-nodulation-cell division (RND) antibiotic efflux pump | antibiotic efflux; antibiotic target alteration | acridine dye; cephalosporin; fluoroquinolone antibiotic; glycylcycline; penam; phenicol antibiotic; rifamycin antibiotic; tetracycline antibiotic; triclosan | fluoroquinolone antibiotic | -1.28 | 10.13 | -4.43 | <0.00001 | 0.0020 |
| MexR | resistance-nodulation-cell division (RND) antibiotic efflux pump | antibiotic efflux; antibiotic target alteration | aminocoumarin antibiotic; carbapenem; cephalosporin; cephamycin; diaminopyrimidine antibiotic; fluoroquinolone antibiotic; macrolide antibiotic; monobactam; penam; penem; peptide antibiotic; phenicol antibiotic; sulfonamide antibiotic; tetracycline antibiotic | fluoroquinolone antibiotic | -0.73 | 11.11 | -3.30 | 0.0015 | 0.0454 |
| MexR | resistance-nodulation-cell division (RND) antibiotic efflux pump | antibiotic efflux; antibiotic target alteration | aminocoumarin antibiotic; carbapenem; cephalosporin; cephamycin; diaminopyrimidine antibiotic; fluoroquinolone antibiotic; macrolide antibiotic; monobactam; penam; penem; peptide antibiotic; phenicol antibiotic; sulfonamide antibiotic; tetracycline antibiotic | monobactam | 1.75 | 10.90 | 4.95 | <0.00001 | 0.0004 |
| mdtO | major facilitator superfamily (MFS) antibiotic efflux pump | antibiotic efflux | acridine dye; nucleoside antibiotic | monobactam | 2.13 | 10.40 | 3.73 | 0.0004 | 0.0150 |
| OpmD | resistance-nodulation-cell division (RND) antibiotic efflux pump | antibiotic efflux | acridine dye; fluoroquinolone antibiotic; tetracycline antibiotic | monobactam | -2.15 | 10.44 | -3.28 | 0.0016 | 0.0414 |
| MexT | resistance-nodulation-cell division (RND) antibiotic efflux pump | antibiotic efflux | diaminopyrimidine antibiotic; fluoroquinolone antibiotic; phenicol antibiotic | monobactam | -1.52 | 10.57 | -3.19 | 0.0021 | 0.0414 |
| MexK | resistance-nodulation-cell division (RND) antibiotic efflux pump | antibiotic efflux | macrolide antibiotic; tetracycline antibiotic; triclosan | nitroimidazole antibiotic | -3.85 | 11.97 | -3.71 | 0.0004 | 0.0407 |

Results of limma analysis were reported in the table together with the genotype, the antibiotic resistance gene name (according to the CARD database), and the antibiotic resistance family (AR family). Other columns are: logFC, the log2-transformed fold change value; AveExpr, the average log2-expression level for that pathway across all samples; t, the t-value according to limma’s moderated t-test; P.Value, the p-value; adj.P.Val, the adjusted p-value using the Benjamini–Hochberg correction. The fold-change value reported refers to the contrast between patients which are treated with an antibiotic sensible to the resistance mechanism specified and patients that were not. Only contrasts with an adjusted p-value lower than 0.05 and an absolute log fold-change value higher than 5 were reported.

**Supplementary Figures**

**FIGURE S1**

Strain-level phylogenetic trees of all detected microbes in the study. Phylogenetic trees obtained through StrainPhlAn pipeline were reported for the main pathogenic signatures of CF disease. Only species with a set of known markers were included in the plot (as reported in the StrainPhlAn pipeline).

**FIGURE S2**

Effect of genotypes and samples on the bacterial diversity of lung microbiome. The effect of a) genotype on alpha diversity was inspected together with b) the interindividual effect. Both the Shannon index and the inverse Simpson index were included in the analysis and reported in different panel. Contrasts reporting a p-value lower than 0.05 were reported using a single asterisk whereas those with a p-value lower than 0.01 were reported using two asterisks. Homozygote and heterozygote refer to ΔF_508_ mutation of CFTR gene. BL, baseline; TR, treatment (samples collected during the treatment of an exacerbation event); RC, recovery (the first sample collected after the end of an exacerbation event).

**FIGURE S3**

Differential abundant pathways. Volcano plot reporting results obtained with limma moderated t-test on pathway distribution are shown. Differential abundant pathways were assessed through the limma moderated t-test and results were reported in this plot. Each panel report a different contrast between samples collected during the treatment of an exacerbation event (TR), after the resolution of an exacerbation event (RC), and during normal visits (BL). Contrasts were divided and grouped according to different genotypes and refers to the ΔF_508_ mutation of CFTR gene. Pathways reporting a significant difference (p-value < 0.05 and |log fold-change| > 5) in the contrast considered were reported in red otherwise were reported in blue. Gray points are those from other contrasts and were reported on the back of the plot.

**FIGURE S4**

Differential abundant antibiotic resistance genes. Volcano plot reporting results obtained with limma moderated t-test on resistance gene distribution are reported. Differential abundant ARGs were assessed through the limma moderated t-test and results were reported in this plot. For additional information about the plot see legend of Figure S3.

**Figure S5**

Effect of the antibiotic intake on the distribution of antibiotic resistance genes. Volcano plot reporting results obtained with limma moderated t-test for each class of antibiotic are shown. Differential abundant tests were performed for each class of antibiotic used in the study. For additional information about the plot see legend of Figure S3.

**Figure S6**

Antibiotic resistance genes map. Antibiotic Resistance Genes (ARGs) were reported in the y-axis whereas samples were reported in the x-axis. Antibiotic classes (both for ARG and for patient treatments) where reported using dots at the end of the heatmap. Hierarchical clustering was computed using the Jaccard index for binary data with the UPGMA method. Red cells correspond to the presence of a gene in samples whereas gray cell correspond to absence. Only genes detected in at least the 10% of the subjects were reported.

**Figure S7**

Antibiotic resistance map of each sample included in the study. Antibiotic resistance genes were reported in the y-axis whereas samples were reported in the x-axis. Antibiotic classes (both for ARG and for patient treatments) where reported using dots at the end of the heatmap. Hierarchical clustering was computed using the Jaccard index for binary data with the UPGMA method. Red cells correspond to the presence of a gene in samples whereas gray cell correspond to absence.
