## Supplementary figures and images for "Taxonomic variability over functional stability in the microbiome of Cystic Fibrosis patients chronically infected by *Pseudomonas aeruginosa*"

### Figure S1

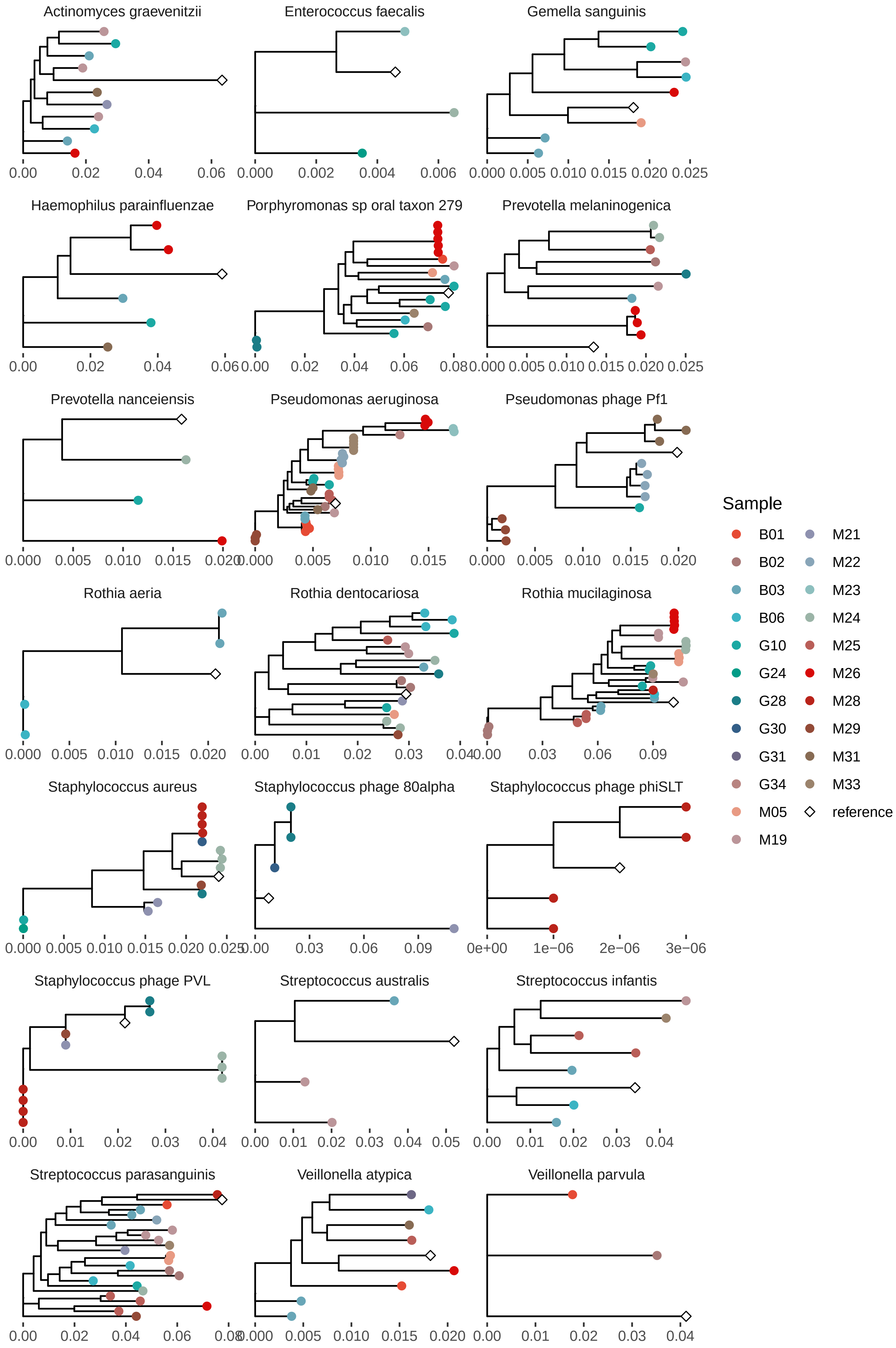

### Figure S2

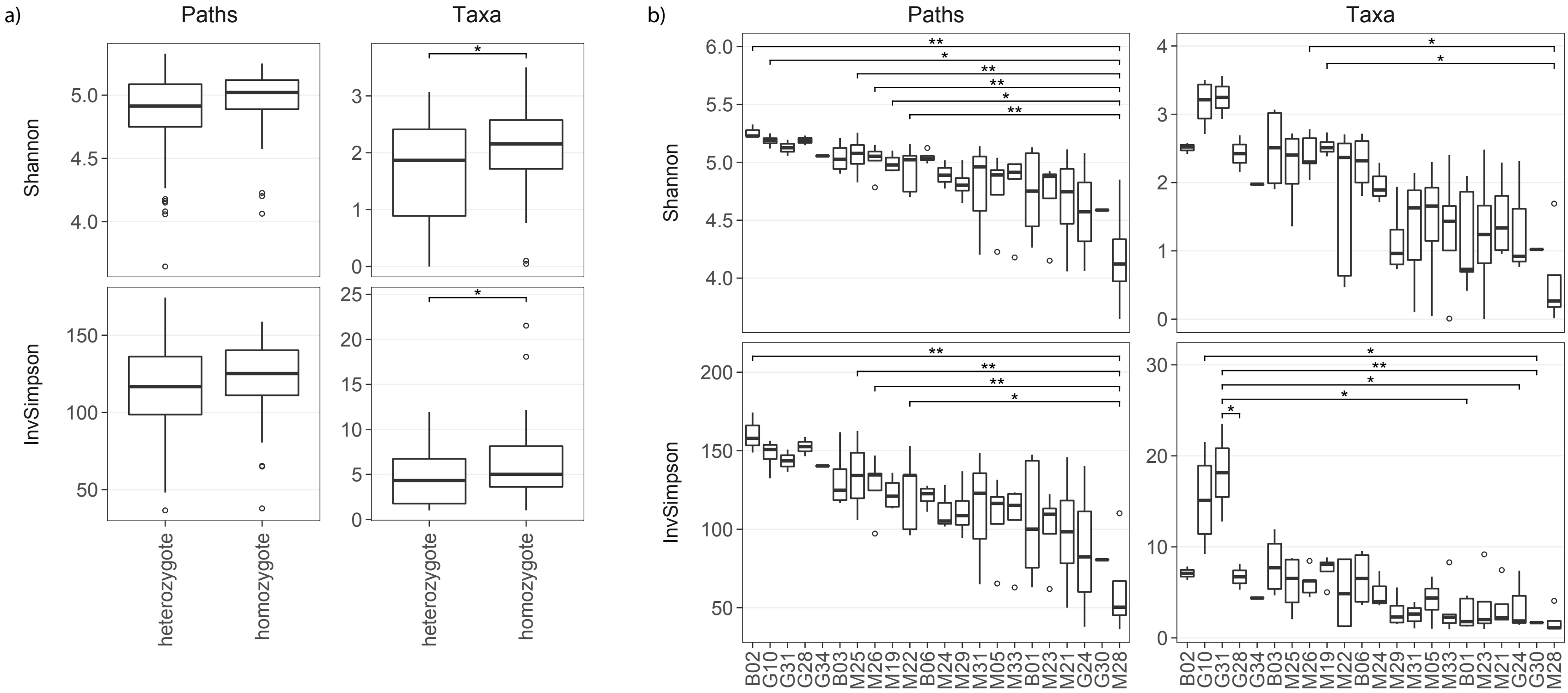

### Figure S3

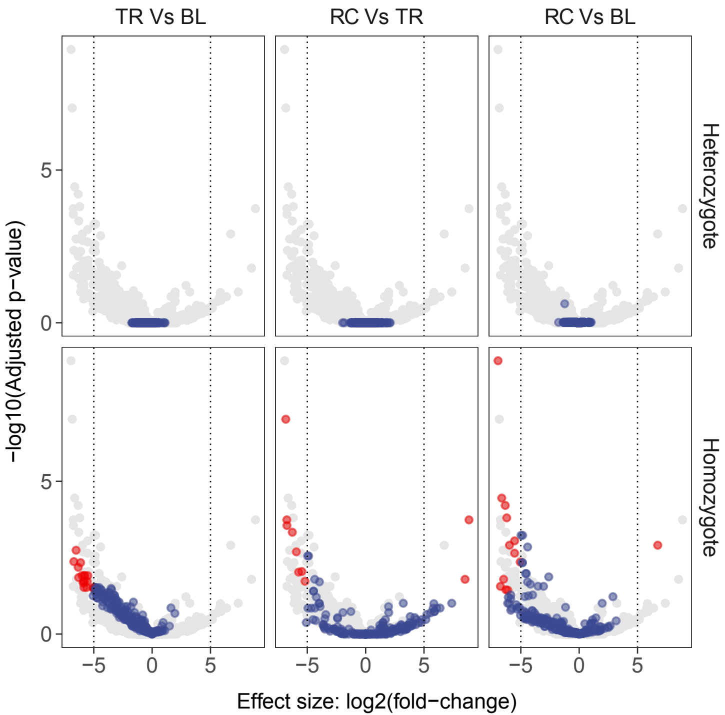

### Figure S4

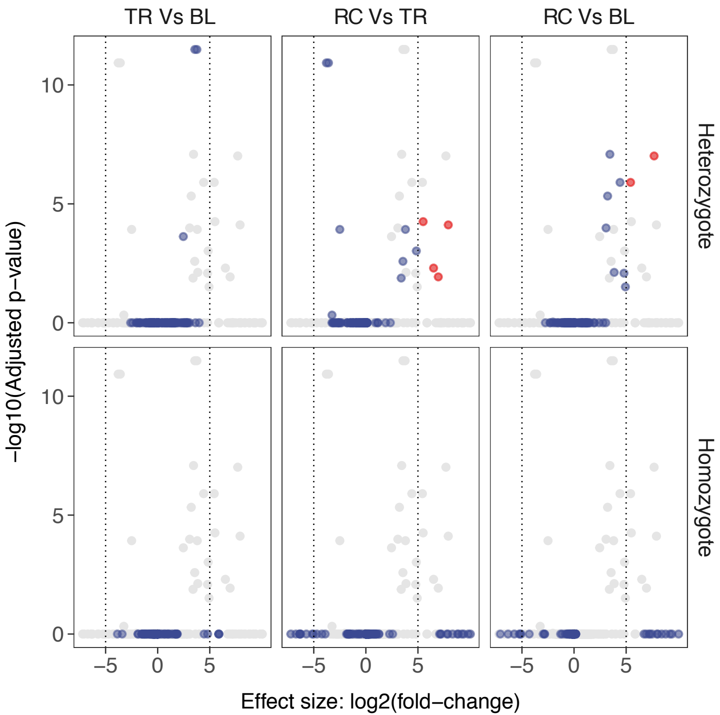

### Figure S5

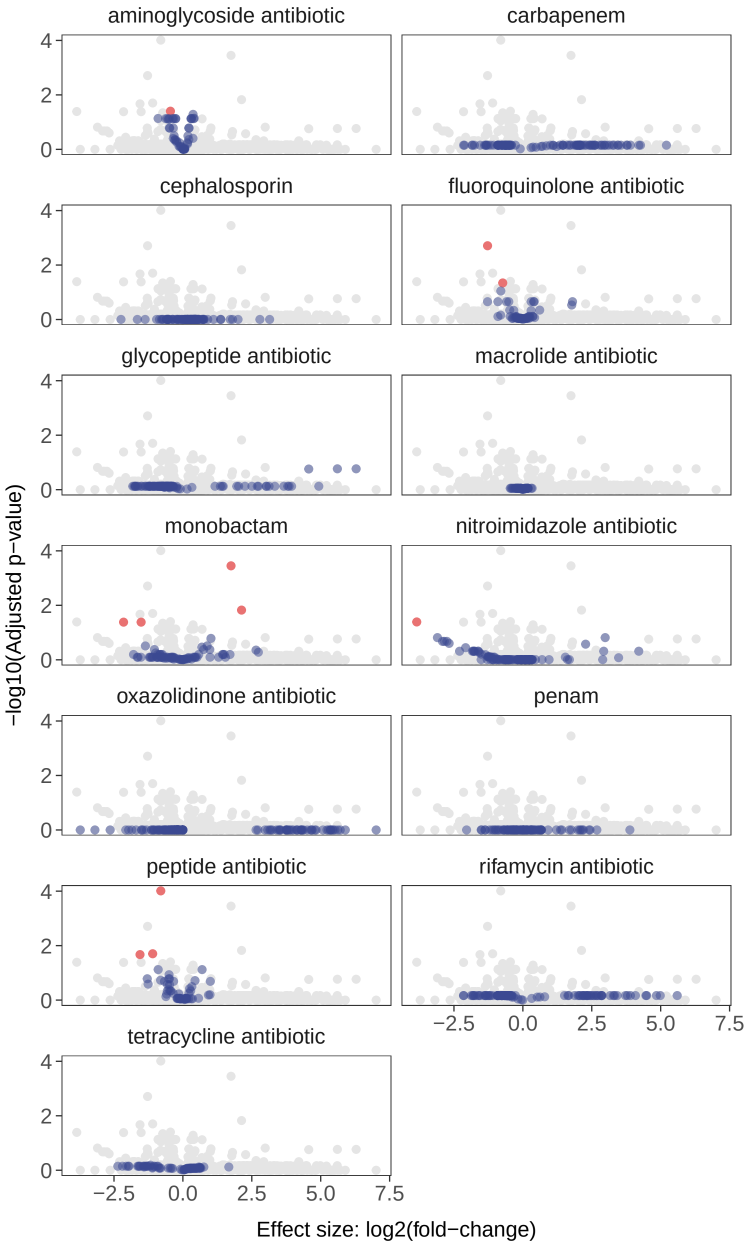

### Figure S6

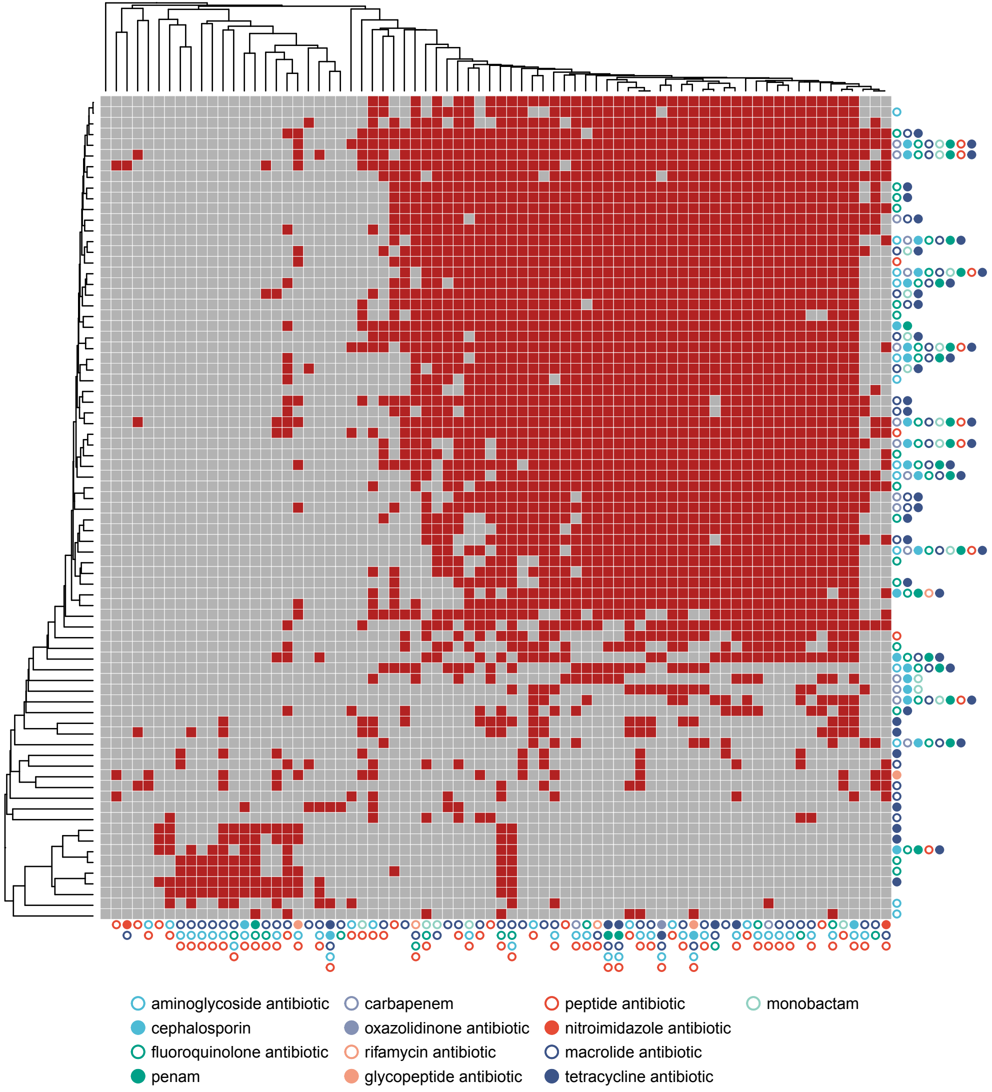

### Figure S7

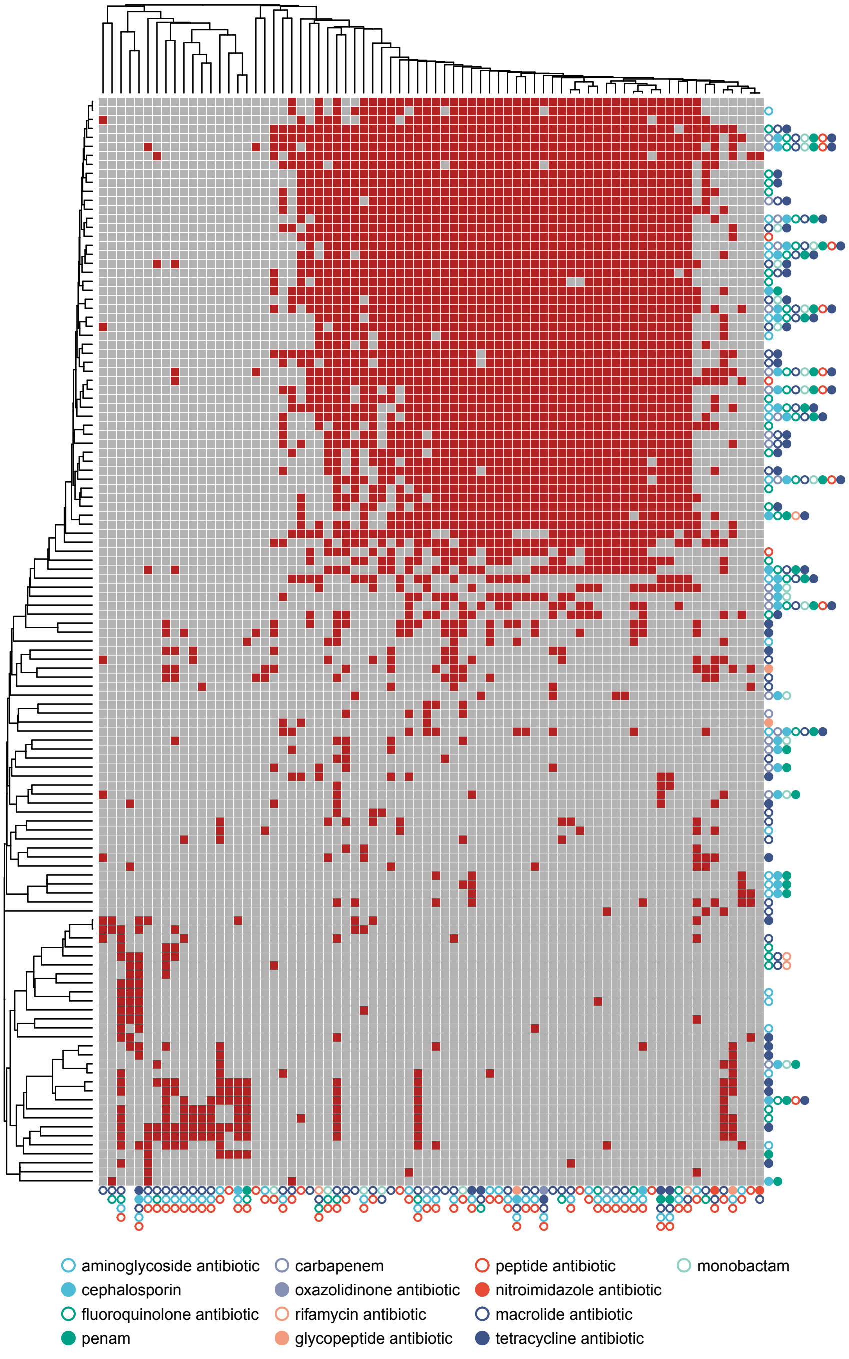
